# A transferrable and integrative type I-F Cascade for heterologous genome editing and transcription modulation

**DOI:** 10.1101/2021.02.08.430362

**Authors:** Zeling Xu, Yanran Li, Huiluo Cao, Meiru Si, Guangming Zhang, Patrick CY Woo, Aixin Yan

**Author notes:** Correspondence (A.Y.).

## Abstract

The Class 1 type I CRISPR-Cas systems represent the most abundant and diverse CRISPR systems in nature. However, their applications for generic genome editing have been hindered by difficulties of introducing the class-specific, multi-component effectors in heterologous hosts for functioning. Here we established a transferrable Cascade system that enables stable integration and expression of a complete and highly active I-F Cascade in the notoriously recalcitrant and diverse *P. aeruginosa* genomes by conjugation. The transferred Cascade displayed substantially higher DNA interference activity and greater editing capacity than the Cas9 system in diverse genetic backgrounds, including removal of the large (21-kb) integrated cassette with efficiency and simplicity. An advanced λred-I-F system enabled editing in genotypes with poor homologous recombination capacity, clinical isolates lacking sequence information, and cells containing anti-CRISPR elements Acrs. Lastly, an ‘all-in-one’ I-F Cascade-mediated CRISPRi platform was developed for transcription modulation by simultaneous introduction of the Cascade and the mini-CRISPR array expressing desired crRNA in one-step. This study provides a framework for expanding the diverse type I Cascades for widespread, heterologous genome editing and establishment of editing techniques in non-model isolates of pathogens.

## INTRODUCTION

Clustered regularly interspaced short palindromic repeats (CRISPR) and CRISPR-associated proteins (Cas) constitute the adaptive immune system in prokaryotes that defends against foreign genetic elements via RNA-guided nucleic acids destruction (1,2). Genome editing and therapeutic applications have focused on the Class 2 CRISPR-Cas systems owing to their simple requirement of a single, multifunctional effector, for example Cas9 and Cas12a, for DNA interference (3,4). However, Class 2 systems represent only ~10% of the CRISPR-Cas systems encoded naturally in prokaryotes (5). Their applications to edit bacterial genomes have been frequently impeded by the poor transformation, cytotoxicity, and requirement of species-specific optimization of the heterologous expression of the large Cas9/Cas12a proteins (6–8). In contrast to the rapid rise and extension of the tools in eukaryotes, thus far, the Cas9/Cas12a-based genome editing is only successfully established in a few model bacterial strains. A predominant, CRISPR-based editing strategy readily applicable in diverse bacterial species is lacking.

Remarkably, nearly 50% of bacteria and 90% of archaea genomes encode native CRISPR-Cas systems and 90% of these naturally occurring CRISPR-Cas systems belongs to the Class 1 systems which target invading nucleic acids via a multi-component effector complex termed as Cascade (CRISPR-associated complex for antiviral defense) (9,10). Although the complexity of these effectors has somewhat hindered their widespread application in eukaryotes, their prevalence and diversity, especially the Class 1 type I systems which account for 50% of all CRISPR-Cas systems identified with seven subtypes (i.e. I-A to I-F plus I-U), has opened new avenues for endogenous CRISPR-Cas-based genetic exploitations in bacteria and archaea (11). The approach operates by simply delivering a programmed mini-CRISPR array and a desired repair donor, which are frequently assembled in a single plasmid, into prokaryotic cells, enabling genome editing with simplicity and efficiency. Employing the strategy, editing of several genetically recalcitrant organisms such as the industrial bacterium *Clostridium pasteurianum* ATCC6013 (type I-B) (12), the multidrug resistant *Pseudomonas aeruginosa* genotype PA154197 (type I-F) (13), and a human microbiome *Lactobacillus crispatus* NCK1350 (Type I-E) (14) has been achieved. Furthermore, the processive DNA degradation fashion of the type I signature nuclease Cas3 has enabled long-range genomic deletions in both bacteria (employing the Type I-C Cascade from *P. aeruginosa*) (15) and human embryonic stem cells (employing the Type I-E Cascade from *T. fusca*) (16) which are inaccessible by the Class 2 effectors. Recently, the type I-B, type I-E, and type I-F systems have also been repurposed for transcription modulation in human cells (17,18). In all these applications, the type I CRISPR-Cas-mediated editing has displayed distinctive advantages, such as high specificity with a minimal off-target effect.

Despite these advantages, currently, the type I CRISPR-Cas-based genome editing is limited to a subset of microbial hosts that harbor a functionally characterized and highly active endogenous CRISPR-Cas system. However, ~ 50% of bacterial genomes are CRISPR-free and a significant portion of natural CRISPR-Cas systems are degenerated without DNA targeting and interference activities (19,20), hindering the widespread application of the endogenous type I Cascade-mediated editing. To overcome this barrier, in this study, we developed a transferrable type I CRISPR-Cas system which enables stable integration and expression of a well-studied, highly active type I-F Cascade in heterologous bacterial hosts for genome editing and transcription modulation. Type I-F Cascade is characterized by the presence of a single DNA-recognizing subunit *cas8f* and the unique *cas2-cas3* gene fusion (Figure 1A) which represents a relatively small size and simple architecture than most other type I systems (21). Furthermore, the subtype occurs exclusively in bacterial organisms, particularly Gammaproteobacteria, underscoring its broad application potentials in this most genera-rich taxa.

**Figure 1.**
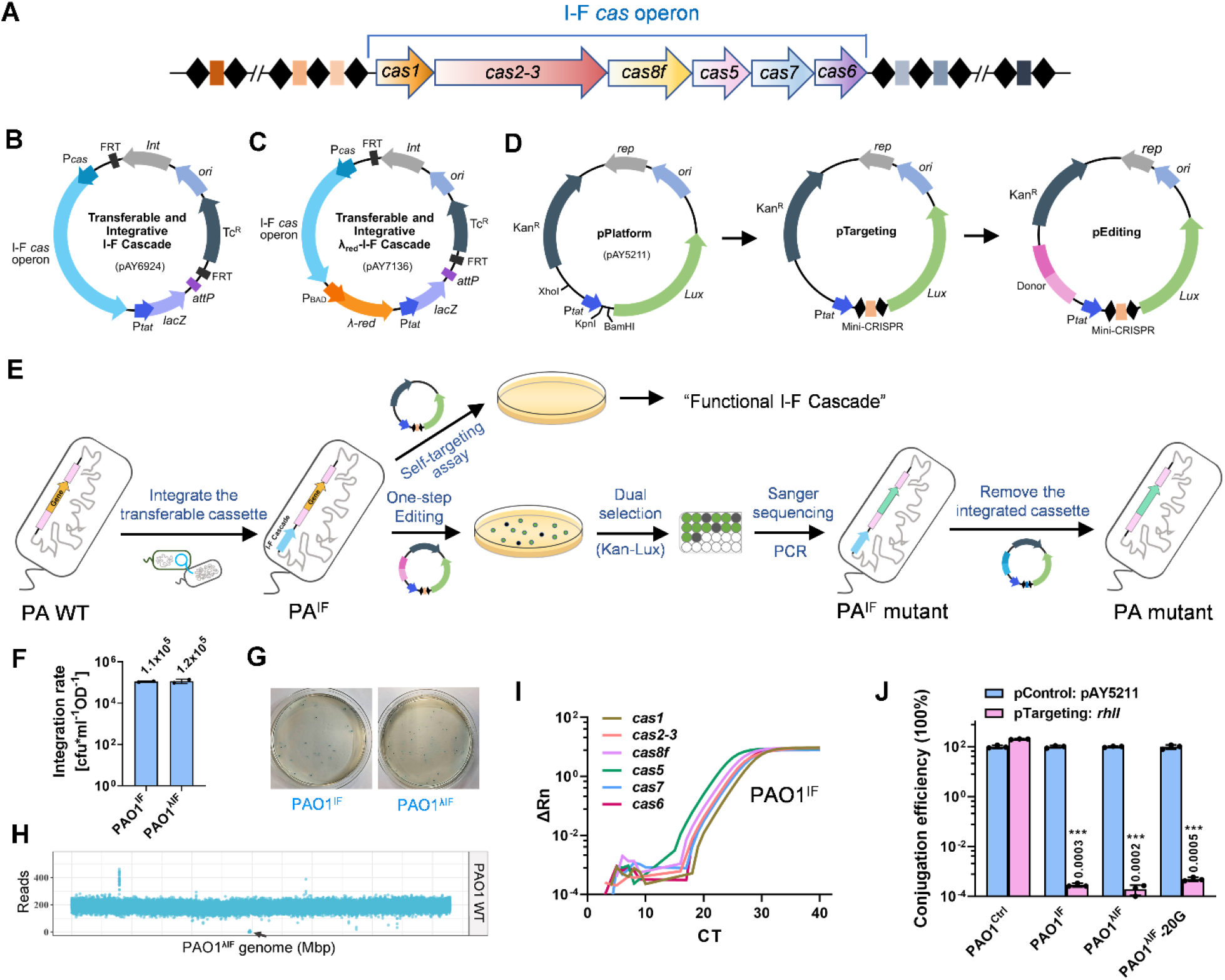
Overview of the transferrable and integrative type I-F Cascade-based genome editing. (**A**) Gene architecture of the type I-F *cas* operon in PA154197. (**B**) Diagram of the transferrable and integrative mini-CTX-IF-P*tat-lacZ* vector (Transferrable I-F Cascade). The organization of genes and elements in the plasmid are shown. The system contains a complete I-F *cas* operon with its native promoter from PA154197, a *lacZ* reporter gene driven by the constitutive promoter P*tat*, a fCTX integrase encoding gene *Int*, a fCTX attachment site *attP* which recognize the chromosomal *attB* site in *P. aeruginosa*, and two Flp recombinase target sites FRT. (**C**) Diagram of the transferrable and integrative λ_red_-I-F Cascade. A phage λ-Red recombination system driven by P_BAD_ is assembled downstream of the I-F *cas* operon. (**D**) Assembly of the targeting plasmid (pTargeting) and editing plasmid (pEditing) for DNA interference assay and gene editing applications, respectively. (**E**) Schematic diagram showing the application of the transferrable I-F Cascade for heterologous genome editing in *P. aeruginosa* (PA). The transferrable I-F Cascade is first integrated into the *attB* site in the genome of a recipient strain (PA WT) to generate a new strain PA^IF^. A targeting plasmid is delivered into PA^IF^ to examine the functionality of the integrated I-F Cascade. An editing plasmid is introduced to achieve desired genome editing. Recovered colonies are inoculated for luminescence-based screening. Luminescence-positive inoculum is subjected to validation by PCR and sanger sequencing. Once the desired editing is achieved, the integrated cassette is removed by another round of editing reaction employing the cassette-deleting editing plasmid (Figure 6A). (**F**) Integration efficiency of the transferrable I-F and λ_red_-I-F Cascade systems. Colony forming unit (CFU) is calculated to indicate the number of cells integrated with the transferrable systems following conjugation of SM10 containing the transferrable systems with PA. (**G**) X-gal-based selection for colonies integrated with the transferrable I-F Cascade. Blue colonies indicate the successful chromosomal integration of the transferrable systems. (**H**) Mapped whole genome sequencing (WGS) reads of the PAO1 wild-type (PAO1 WT) strain against the PAO1^λIF^ genome. Arrow indicates the absence of reads of the integrated cassette in PAO1 WT at the *attB* site compared with PAO1^λIF^. (**I**) Quantitative PCR analysis of the expression of the integrated *cas* genes in PAO1^IF^. Rn denotes the reporter signal normalized to the fluorescence signal of the passive reference dye (ROX). ΔRn is the value of Rn minus the baseline. (**J**) Self-targeting assay as indicated by the conjugation efficiency following introduction of the targeting and control plasmids in the indicated strains. PAO1^Ctrl^: PAO1 strain integrated with the empty transferrable system lacking the *cas* operon; PAO1^IF^: PAO1 strain integrated with the transferrable I-F Cascade; PAO1^λIF^: PAO1 strain integrated with the transferrable λ_red_-I-F Cascade; PAO1^λIF^-20G: the PAO1^λIF^ strain after growing 20 generations in the absence of antibiotic selection. Data are displayed as mean ± S.D from three biological repeats. Statistical significance is calculated based on student’s *t* test (***, *P* < 0.001).

The Gammaproteobacterium *Pseudomonas aeruginosa* is a primary model organism for the type I-F CRISPR-Cas system. The pathogen is particularly notorious for its diverse genotypes (22), rendering difficulty of establishing genetic tools in the strains of interests with clinical, environmental, or industrial significance. Hence, we establish the transferrable I-F system and test its robustness using the genetically recalcitrant gammaproteobacterium *P. aeruginosa* as a paradigm. Employing the integration-proficient transferrable I-F system mini-CTX-IF-*lacZ* we developed, we show that the highly active type I-F Cascade from the PA154197 strain is stably integrated and expressed in diverse *P. aeruginosa* genotypes, enabling endogenous CRISPR-Cas-based genetic exploitations in the transferred hosts. Furthermore, an advanced mini-CTX-λ_red_-IF-*lacZ* system is developed to achieve genome editing in homologous recombination poor genotypes, including strains lacking genome sequence information. Following the desired genome editing, the integrated cassette (21.212 kb) encompassing the entire I-F Cascade genes, a *lacZ* reporter, and a λ-Red recombination system is readily removed by employing a common cassette-deleting editing plasmid we devised. We also show that the system is applicable in strains containing anti-CRISPR (Acr) elements following firstly removing *acrs* by the counter selection-based approach. Lastly, a I-F CRISPRi platform which lacks the nuclease Cas2-3 and is equipped with a multiple cloning site for incorporation of a programmed mini-CRISPR array enabled one-step gene transcription modulation in heterologous hosts. This approach opens a new avenue for widespread exploitation of the diverse type I CRISPR-Cas systems for heterologous genome editing.

## MATERIALS AND METHODS

### Plasmids, primers, and bacterial growth conditions

All the bacterial strains, plasmids and primers used in this study are summarized in the Supplementary Table S1. *E. coli* DH5α was employed for plasmid propagation and was usually cultured at 37°C in Luria-Bertani (LB) broth or on the LB agar plate supplemented with required antibiotics. *E. coli* SM10 was employed for conjugative plasmid delivery. *P. aeruginosa* clinical strains were isolated from the Queen Mary Hospital in Hong Kong, China. Antibiotics supplemented in the agar plates for culturing DH5α were 20 μg/ml kanamycin, 10 μg/ml tetracycline or 200 μg/ml ampicillin. Antibiotics supplemented in the agar plates for culturing SM10 were 500 μg/ml kanamycin, 10 μg/ml tetracycline or 200 μg/ml ampicillin. Antibiotics supplemented in the agar plates for culturing *P. aeruginosa* strains were 500 μg/ml kanamycin, 50 μg/ml tetracycline or 200 μg/ml carbenicillin.

### Construction of the mini-CTX-IF-P*tat-lacZ* and mini-CTX-λ_red_-IF-P*tat-lacZ*

The transferrable I-F Cascade system was constructed based on the mini-CTX-*lacZ* plasmid. The mini-CTX-*lacZ* plasmid was linearized by HindIII (NEB, USA) at 37°C for 4 hrs. The P*tat* promoter DNA fragment was obtained by PCR using the iProof High-Fidelity DNA Polymerase (Bio-Rad, USA) and PA154197 genomic DNA as the template. The fragment was ligated into the HindIII-digested mini-CTX-*lacZ* plasmid following the instruction of the ClonExpress II One Step Cloning Kit (Vazyme, China), generating mini-CTX-P*tat-lacZ*. The I-F *cas* operon including its native promoter (8.693-kb) was obtained by PCR employing PA154197 genomic DNA as the template. The DNA fragment was inserted into the KpnI site in the mini-CTX-P*tat-lacZ* which was digested with KpnI (NEB, USA) using the ClonExpress II One Step Cloning Kit, generating a new plasmid mini-CTX-IF-P*tat-lacZ* (Transferrable I-F Cascade, pAY6924). To construct the transferrable λ_red_-I-F Cascade, the fragment containing the λ-Red genes and the L-arabinose inducible P_BAD_ promoter was first obtained by PCR employing pKD46 plasmid as the template. The fragment was inserted into the SalI sites in the mini-CTX-IF-P*tat-lacZ* using the ClonExpress II One Step Cloning Kit, generating the new plasmid mini-CTX-λ_red_-IF-P*tat-lacZ* (Transferrable λ_red_-I-F Cascade, pAY7136). To construct the transferrable λ_red_-Cas9 system, the I-F *cas* operon and its promoter in the transferrable λ_red_-I-F Cascade was replaced by the Sp*cas9* gene with the P_BAD_ promoter which was amplified from pCasPA (23).

### Integration of the transferrable I-F Cascade

*E. coli* SM10 strain transformed with the plasmid pAY6924 (transferrable I-F Cascade) was cultured in LB broth supplemented with 10 μg/ml tetracycline at 37°C with 220-rpm agitation for 16 hrs. Simultaneously, *P. aeruginosa* recipient strain was cultured in LB broth at 42°C with 220-rpm agitation for 16 hrs. Cell densities of the *E. coli* SM10 and *P. aeruginosa* cultures were determined by measuring OD_600_. Subsequently, 1.5×10^9^ *E. coli* SM10 and 0.25×10^9^ *P. aeruginosa* cells were mixed and pelleted by centrifugation at 16,000 ×g for 1 min. Cell pellet was resuspended in 50 μl LB broth and subsequently spotted on the surface of a LB agar plate. Mating (plasmid delivery from SM10 to *P. aeruginosa*) occurred during the incubation of the mixture at 37°C for 8 hrs. Mixture was scrapped from the agar plate and resuspended in 300 μl PBS (Phosphate Buffered Saline, pH 7.4). Cell suspension was serial diluted and spread on the VBMM plates with supplementation of 50 μg/ml tetracycline and 40 μg/ml X-gal. Plates were incubated at 37°C for 24 hrs. Blue colonies indicating the chromosomal integration of the transferrable system were selected for further analysis.

### Reverse transcription-quantitative PCR (RT-qPCR)

Expression of the integrated genes such as *cas* and λ-Red genes was examined by RT-qPCR following previous descriptions (24). Briefly, 1 ml bacterial cells cultured in LB were harvested when the OD600 reached 1.0. Total RNA was extracted employing the Takara MiniBEST Universal RNA Extraction Kit (Takara, Japan) and reverse transcription was conducted employing the PrimeScript RT Master Mix (Takara, Japan) following the manufacturer’s instructions. qPCR was performed using specific primers mixed with the TB Green Premix Ex Taq (Takara, Japan) in a 20 μl reaction system. The amplification was performed in the ABI StepOnePlus real-time PCR system. Amplification curves were plotted to show the transcription levels of the genes of interest. For relative quantification of gene expression, 2^-ΔΔCt^ method (25) was applied and the *recA* gene was selected as the internal reference gene.

### Construction of pTargeting and pEditing

A platform plasmid (pAY-mini-CRISPR, pAY6942) was firstly constructed to facilitate the construction of the desired mini-CRISPR elements required in various editing procedures. pAY6942 encompasses an insertion site encompassed by a tandem BsaI sites which are flanked by a repeat sequence (GTTCACTGCCGTATAGGCAGCTAAGAAA) specific to the I-F system at each end. A fragment of 32-bp nucleotides (<u>N×32</u>) preceded by a 5’-CC-3’ PAM in the coding region of the gene to be edited was firstly identified as the protospacer. Two 36-bp oligos encompassing the selected protospacer sequence flanked by four nucleotides overlapping with the 3’-end and 5’-end of the repeat sequences, i.e. 5’-GAAA<u>N×32</u>-3’ and 5’-GAAC<u>N×32</u>-3’, respectively, were synthesized commercially (IDT, Singapore). The two oligos were then phosphorylated using T4 polynucleotide kinase (NEB, USA) at 37°C for 1 hr. The phosphorylated oligos were heated at 95°C for 3 min and then cooled down to room temperature to generate annealed oligos. Annealed oligos were ligated into the BsaI-digested pAY6942 using the Quick LigationTM Kit (NEB, USA), resulting in a plasmid containing the desired mini-CRISPR array. Assembly of mini-CRISPR into the platform plasmid (pAY5211) results in the self-targeting plasmid (pTargeting). Further assembly of the donor template into pTargeting results in pEditing. The procedures were described previously (26). Briefly, amplified mini-CRISPR elements were digested by KpnI and BamHI (NEB, USA) and were ligated into pAY5211 employing the Quick LigationTM Kit (NEB, USA), resulting in the targeting plasmid (pTargeting). Donor sequences which usually contain the upstream and downstream homologous arms of the gene being edited with 21-bp overlap of the XhoI-digested targeting plasmid at each end were amplified by PCR and ligated into the linearized targeting plasmid (digested by XhoI (NEB, USA)) using the ClonExpress II One Step Cloning Kit (Vazyme, China), resulting in the editing plasmid (pEditing). All plasmids constructed were verified by Sanger sequencing (BGI, China).

### Quantification of conjugation efficiency

*E. coli* SM10 strain harboring the plasmid pAY5211 or pAY7138 was cultured in LB broth supplemented with 100 μg/ml kanamycin at 37°C with 220-rpm agitation for 16 hrs. Simultaneously, *P. aeruginosa* strains containing the I-F or λ_red_-I-F Cascade were cultured in LB broth at 42°C with 220-rpm agitation for 16 hrs. Conjugation of *E. coli* SM10 with *P. aeruginosa* was performed as described above. After conjugation, mixture was scrapped from the agar plate and resuspended in 300 μl PBS buffer. The cell suspension was subsequently adjusted to OD_600_ as 1.0 and was serially diluted and plated on the LB agar plates containing 50 μg/ml tetracycline and 500 μg/ml kanamycin (tetracycline resistance of the *P. aeruginosa* recipients due to the integrated transferrable I-F Cascade). Plates were incubated at 37°C for 24-36 hrs. Numbers of recovered cells from the conjugation reaction was determined by CFU obtained multiply the dilution factors. Colony number recovered from pAY5211 was set as 100%.

### Genome editing and verification

*E. coli* SM10 strain transformed with the editing plasmid for desired gene editing was cultured in LB broth supplemented with 100 μg/ml kanamycin at 37°C with 220-rpm agitation for 16 hrs. *P. aeruginosa* recipient strains integrated with the λ_red_-I-F Cascade were cultured in LB broth supplemented with 20 mM L-arabinose at 42°C with 220-rpm agitation for 16 hrs. Conjugation of *E. coli* SM10 and *P. aeruginosa* were performed as described above. After conjugation, the cell mixture was scrapped from the agar plate and resuspended in 300 μl PBS buffer. 150 μl cell suspension was spread on the LB agar plates containing 50 μg/ml tetracycline and 500 μg/ml kanamycin. Plates were incubated at 37°C for 24-36 hrs. Recovered single colonies were inoculated into LB broth containing 100 μg/ml kanamycin in a 96-well plate. After incubation at 37°C with 220-rpm agitation for 3 h, luminescence intensity was measured and ten colonies with the highest luminescence intensity were subjected to verification by colony PCR and sanger sequencing with specific primers. The editing plasmid (pEditing) in the edited *P. aeruginosa* cells was cured by streaking the cells onto a LB ager plate and incubation at 37°C for 16 hrs. In certain genotypes, multiple (2 to 3) rounds of streaking are required for thorough plasmid curing.

### Bacterial whole genome sequencing and analysis

Genomic DNA was extracted from 1 ml overnight bacterial culture using the Illustra bacteria genomicPrep Mini Spin Kit (GE Healthcare, USA) according to the manufacturer’s instructions. Whole genome sequencing was conducted by Novogene (Beijing, China) with the sequencing platform Novaseq. Raw reads were subjected to quality control using Trimmomatic(27). The genome of the PAO1^λIF^ strain was assembled using SPAdes with PAO1 (NC_002516.2) as the reference (28) and completed using an assembly improvement pipeline (29). Circlator’s fixstart task was applied to fix the start position of the manually assembled complete genome of PAO1^λIF^ at the *dnaA* gene (30). Annotations of PAO1^λIF^ were conducted using Prokka with *Pseudomonas* genera specific database (31). Genome-wide integration site distribution for PAO1^λIF^ was conducted with the complete genome of PAO1 as the reference following previous descriptions (32). Mapping in the present study was performed using BWA (33). Integration events within 500 bp bins were computed using SAMtools (34) and BEDTools (35) and visualized with ggplot2 in R platform (https://ggplot2.tidyverse.org/index.html). To explore mutations in the three PAO1^λIF^ *ΔrhlI* strains and three false positive strains relative to the PAO1^λIF^, core SNPs were collected and annotated with the PAO1^λIF^ genome as reference using snippy (https://github.com/tseemann/snippy).

### PYO quantification

PYO production was quantified following the protocol described previously with slight modifications (36). 1 ml bacterial culture was subjected to centrifugation at 16,000 ×g for 5 min. 750 μl supernatant was collected and mixed with 450 μl chloroform by vortex for 0.5 min. After centrifugation at 16,000 ×g for 5 min, 400 μl liquid from lower phase was then mixed with 200 μl HCl (0.2 M) by vortex thoroughly for 0.5 min. After centrifugation at 16,000 ×g for 5 min, 100 μl upper aqueous phase containing PYO was transferred to a 96-well plate and its absorbance was measured at 510 nm. Concentration of PYO was determined according to the standard curve.

## RESULTS

### A transferrable and integrative I-F Cascade is developed and is active in the recipient *P. aeruginosa* host

Our previous study has demonstrated that the type I-F CRISPR-Cas system encoded in the *P. aeruginosa* PA154197 strain is highly active for DNA interference and genome editing (13). Hence, we set out to construct a transferrable I-F Cascade for heterologous applications based on this system. The I-F Cascade encompasses Cas8f (Csy1), Cas5 (Csy2), Cas7 (Csy3) and Cas6 (Csy4) proteins which constitute the complex for CRISPR RNA (crRNA) biogenesis and DNA targeting and the Cas2-3 fusion protein for DNA degradation (Figure 1A) (37,38). To enable stable expression of a heterologous Cascade in recipient cells and to circumvent the dependence of antibiotics to maintain the system, we employ an integration-proficient vector containing a *lacZ* reporter, mini-CTX-*lacZ* (39), to deliver the I-F Cascade. The vector carries an fCTX integrase gene *Int* and a single fCTX attachment site *attP* which enable the integration of the entire backbone into the highly conserved *attB* site in *P. aeruginosa* genomes *via* site-specific recombination (Figure 1B). The complete I-F *cas* operon with its native promoter (8.693 kb) was amplified from PA154197 genomic DNA and was cloned to the KpnI site in the mini-CTX-*lacZ* vector downstream of the FRT site for integration. To facilitate the screening of integration-positive clones, a strong housekeeping promoter P*tat* (40) is placed upstream of the promoter-less *lacZ* in the original vector for blue/white colony selection, resulting in a 17.757-kb mini-CTX-IF-*Ptat-lacZ* plasmid (termed as the transferrable I-F Cascade, pAY6924) (Figure 1B). Considering that genetic modifications in prokaryotes predominantly rely on the homology-directed repair (HDR) and most bacteria lack an efficient intrinsic homologous recombination system, we also developed an advanced transferrable I-F plasmid to simultaneously include an inducible phage λ-Red recombination system (AraC, P_BAD_, Exo, Gam and Beta, 3.455 kb) at the SalI site downstream of the I-F *cas* operon, yielding a 21.212-kb transferrable plasmid mini-CTX-λ_red_-IF-P*tat-lacZ* termed as the transferrable λ_red_-I-F Cascade (pAY7136) (Figure 1C). Integration of these transferrable I-F Cascade systems should allow a *P. aeruginosa* host to temporarily acquire a ‘native’ I-F *cas* operon in its genome, enabling the exploitation of native CRISPR-Cas-based genome editing which is efficient and simple (Figure 1D and 1E) (12–14).

To examine the transferability and applicability of the system, we first delivered the transferrable I-F Cascade (pAY6924) and the transferrable λ_red_-I-F Cascade (pAY7136) into a CRISPR-free, model strain PAO1 by conjugation. 1.1×10^5^ and 1.2×10^5^ clones were recovered per conjugation (Figure 1F), respectively, and all the clones displayed blue color (Figure 1G), suggesting the successful and efficient chromosomal integration of the cassettes. The resulting constructs were termed as PAO1^IF^ and PAO1^λIF^, respectively. A representative PAO1^λIF^ clone was subjected to whole genome sequencing (WGS) which revealed that the cassette was site-specifically integrated at the *attB* site (Genomic location: 2,947,580-2,947,610) (Figure 1H) and no undesirable genetic changes occurred except for a 4-bp synonymous substitution in the *ccoN1_3* gene (Supplementary Table S2). RT-qPCR analysis showed that the integrated *cas* genes were expressed in stoichiometry in the transferred host PAO1^IF^ (Figure 1I) and did not cause alterations in the physiology of the host (Supplementary Figure S1). Whole genome transcriptome analysis and RT-qPCR quantification identified only one differentially expressed gene PA1137 (2-fold higher) encoding a predicted oxidoreductase in the transferred host PAO1^IF^ relative to the PAO1 WT (Supplementary Figure S1B and S1C). These results demonstrated that the integration and expression of the I-F Cascade does not perturb the physiology of the host strain.

Next, we assessed the DNA interference activity of the transferred I-F Cascade by supplying a self-targeting plasmid (pAY7181) which expresses a crRNA targeting to a genomic locus *rhlI* gene (11). Delivery of the plasmid into PAO1^IF^ and PAO1^λIF^ by conjugation resulted in a conjugation recovery rate 6-magnitude lower (conjugation efficiency of 0.002% and 0.003%, respectively) than that of the control plasmid pAY5211 (Figure 1J), indicating that the transferred I-F Cascade displays a high DNA interference activity in the transferred hosts PAO1^IF^ and PAO1^λIF^, resulting in genome cleavage and cell death. The high DNA interference activity was maintained in the PAO1^λIF^ host after 20 generations of growth in the absence of antibiotic selection (Figure 1J), demonstrating the stable integration and function of the transferred I-F Cascade in the host strain.

### Transferrable type I-F system displays substantially greater DNA interference capacity than the transferrable Cas9 system

We next compared the DNA interference activity of the transferrable type I-F Cascade we developed with a transferrable Cas9 system. A similar transferrable Cas9 system (pAY7166) was constructed by replacing the I-F Cascade and its promoter in the mini-CTX-λ_red_-IF-P*tat-lacZ* vector (pAY7136) with the Sp*cas9* gene driven by a P_BAD_ promoter (Supplementary Figure S2A). Integration of pAY7166 into PAO1 generated a PAO1 variant integrated with the Sp*cas9* and λ-Red genes at the *attB* site, termed as PAO1^λCas9^. The expression of *cas9* and λ-Red genes in this strain was confirmed by RT-qPCR analysis (Supplementary Figure S2B). Considering that the type I-F Cascade recognizes a PAM sequence of 5’-CC-3’ located at the 5’-end of a 32-bp protospacer and the Cas9 effector recognizes a PAM sequence of 5’-NGG-3’ located at the 3’-end of a 20-bp protospacer (41), to compare the DNA interference activity of the transferred I-F and Cas9 systems, we constructed pairs of self-targeting plasmids (pTargeting) that express crRNAs recognized by the two effectors, respectively, and target to protospacers with maximal overlap (Figure 2A). These three pairs of protospacers are located in the integrated region, including one pair in the P*tat* promoter (IF_Ps1/Cas9_Ps1) and two pairs in the *lacZ* gene (IF_Ps2/ Cas9_Ps2 and IF_Ps3/Cas9_Ps3) (Figure 2A). Following delivery of each of the targeting plasmids into PAO1^λIF^ or PAO1^λCas9^ by conjugation, the numbers of recovered colonies relative to that of the corresponding control plasmid which does not contain the programmed self-targeting mini-CRISPR are recorded to indicate the DNA interference activity. The transferred I-F Cascade effected a strong self-targeting activity toward all three selected protospacers as evidenced by a 6-magnitude reduction of colony recovery relative to that of the control plasmid, i.e. conjugation efficiency of 0.0007% (IF_Ps1), 0.0003% (IF_Ps2), and 0.0001% (IF_Ps3) (Figure 2C), respectively. However, the transferred Cas9 system only effected 1-magnitude reduction of colony recovery upon introduction of the three pTargeting plasmids relative to the control plasmid, i.e. conjugation efficiency of 25.2% (Cas9_Ps1), 45.6% (Cas9_Ps2) and 52.0% (Cas9_Ps3), respectively (Figure 2D), indicating that the transferred I-F Cascade displays a substantially greater targeting capacity than the transferred Cas9 system.

**Figure 2.**
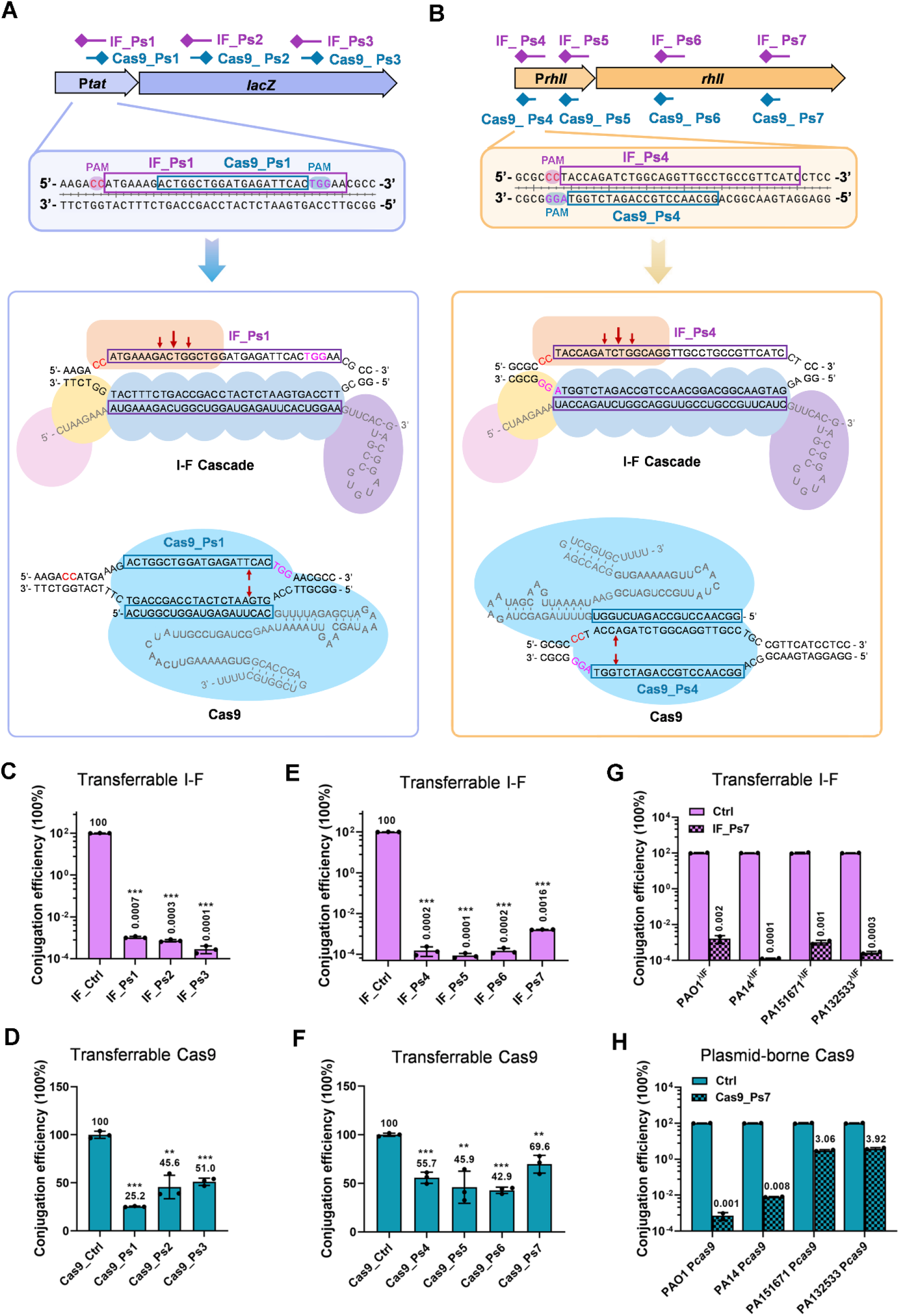
The transferrable I-F Cascade displays substantially greater targeting capacity than the transferrable and the plasmid-borne Cas9 systems. (**A, B**) Schematic diagram of the DNA targeting assay. Pairs of the targeting sites (protospacers) located in both the integrated region *Ptat-lacZ* (A) and the endogenous region *PrhlI-rhlI* (B) were employed to compare the DNA interference activity of the transferred type I-F Cascade and the transferred Cas9 in PAO1^λIF^ and PAO1^λCas9^ cells, respectively. Each pair of the protospacers contains one applied for the I-F Cascade and one for Cas9 with following features: three pairs (IF_Ps1/Cas9_Ps1, IF_Ps2/Cas9_Ps2 and IF_Ps3/Cas9_Ps3) are located in the same DNA strand with maximal overlap in each pair (A); four pairs (IF_Ps4/Cas9_Ps4, IF_Ps5/Cas9_Ps5, IF_Ps6/Cas9_Ps6 and IF_Ps7/Cas9_Ps7) are located in the complementary strands sharing the same PAM location (B). Diamonds denote the PAM sequences. The nucleotide sequences of the protospacer pair IF_Ps1/Cas9_Ps1 (A) and IF_Ps4/Cas9_Ps4 (B) and the resulting targeting pattens mediated by the I-F Cascade and Cas9 are shown in the lower panels. The 32-bp protospacer for the I-F Cascade targeting is framed in purple boxes and the 20-bp protospacer for Cas9 targeting is framed in blue boxes. (**C**, **D**) DNA interference activity of the transferred I-F Cascade (C) or the transferred Cas9 effector (D) determined by the colony recovery rate of pTargeting transformation relative to control plasmid (lacking the mini-CRISPR array) transformation via conjugation, i.e. conjugation efficiency. The protospacer site targeted by the employed pTargeting is shown (x-axis). A low conjugation efficiency indicates a high DNA interference activity exhibited by the effector examined. (**E**, **F**) DNA interference activity of the transferred I-F Cascade (E) or the transferred Cas9 (F) determined by the conjugation efficiency of indicated pTargeting transformation. pTargeting is denoted by the corresponding protospacer site (x-axis) it targets to. (**G, H**) DNA interference activity of the transferred I-F Cascade (G) or the plasmid-borne Cas9 (H) determined by the conjugation efficiency of transforming the pTargeting plasmids that target to the IF_Ps7 (G) or Cas9_Ps7 (H) site, respectively, into the indicated host strains PAO1, PA14, PA151671 and PA132533 (x-axis). Data are the mean of three (B, C, D, F) or two (G, H) biological repeats and are expressed as mean ± S.D. Statistical significance is calculated based on student’s *t* test (*, *P* < 0.05; **, *P* < 0.01; ***, *P* < 0.001).

To exclude the possibility that the poor targeting activity of the transferred Cas9 was due to the locations of the protospacers selected, we examined the targeting of the two effectors to four additional pairs of protospacers located in an endogenous gene locus *rhlI*. The four pairs of spacer sequences employed share an identical PAM location distributed in the complementary strand of the protospacer and display 20-bp complementarity (Figure 2B). Following introduction of the corresponding pTargeting into PAO1^λIF^ and PAO1^λCas9^, the transferred I-F system displayed a substantially higher targeting capacity to all these four sites than the transferred Cas9 system, as evidenced by the conjugation efficiency of 0.0002% (IF_Ps4), 0.0001% (IF_Ps5), 0.0002% (IF_Ps6) and 0.0016% (IF_Ps7) effected by the I-F system as opposed to 55.7% (Cas9_Ps4), 45.9% (Cas9_Ps5),42.9% (Cas9_Ps6) and 69.6% (Cas9_Ps7) effected by the Cas9 system (Figure 2E and 2F), further demonstrating the substantially higher targeting capacity of the transferred I-F system than the Cas9 system in *P. aeruginosa*.

Activity of the transferrable I-F Cascade was further compared with a plasmid-borne Cas9 which was developed and employed by Chen *et al* in a modified two-plasmid-based genome-editing system (23). A pair of pTargeting that targets to the protospacer IF_Ps7 and Cas9_Ps7 located in the *rhlI* gene was employed (Figure 2B). The transferred I-F Cascade was found to display a comparable DNA interference activity with this plasmid-borne Cas9 as demonstrated by the similar conjugation efficiencies (0.001-0.002%) effected by the two effectors (Figure 2G and 2H). However, when these two systems were assessed in several additional *P. aeruginosa* strains such as PA14 and two clinical isolates PA151671 and PA132533, we found that the transferrable I-F Cascade displayed a greater targeting capacity than the plasmid-borne Cas9 as demonstrated by the conjugation efficiency of the pTargeting (IF_Ps7) as 0.0001%, 0.001% and 0.0003%, respectively, in PA14^λIF^, PA151671^λIF^ and PA132533^λIF^, whereas that of the corresponding pTargeting (Cas9_IF7) effected by the plasmid-borne Cas9 system were 0.008%, 3.06% and 3.92%, respectively (Figure 2G and 2H). These results demonstrated that the transferrable I-F Cascade system exhibits substantially greater DNA interference capacity and efficiency than both the transferrable and plasmid-borne Cas9-based systems, underscoring its application potentials for widespread heterologous genome editing.

### Transferred I-F Cascade is reprogrammable for various genetic exploitations in the model heterologous host PAO1

We next examined the applicability of the transferred I-F Cascade in PAO1^IF^ for genome editing using deleting *rhlI* gene as an example which has been shown to display a readily detectable phenotype, i.e. loss of the green-blue pigmented pyocyanin (PYO) production compared with the WT (42). We employed the strategy established for the native I-F Cascade meditated genome editing which is achieved by one-step transformation of a single editing plasmid (13). An editing plasmid termed as p*rhlI*-Del-1 (pAY7420) was constructed by assembling ~800-bp upstream and ~800-bp downstream homologous arms of *rhlI* into the *rhlI*-targeting plasmid pAY7181 employed in the self-targeting assay for HR-mediated *rhlI* deletion (Figure 3A and Supplementary Figure S3A) (26). The editing plasmid also encodes a *lux* operon in its backbone (13), enabling luminescence-based screening for the desired colonies following introduction of p*rhlI*-Del-1 into PAO1^IF^ by conjugation. Ten clones displaying high luminescence intensity were analyzed by colony PCR and all clones were shown to harbor the desired *rhlI* deletion (Figure 3B and Supplementary Figure S3B). This result demonstrated the applicability and high efficiency of the transferred type I-F Cascade for genome editing in the recipient host. To evaluate the editing efficiency in a larger population of recovered clones, we examined 48 luminescence-positive clones by colony PCR which showed that 37 clones harbored the desired *rhlI* deletion (Supplementary Figure S3C). Abolishment of the green-blue pigmented PYO was also observed in all 37 clones compared to the WT PAO1 (Supplementary Figure S3D). An average efficiency of 81.3% was obtained based on two independently conducted editing reactions (Figure 3C).

**Figure 3.**
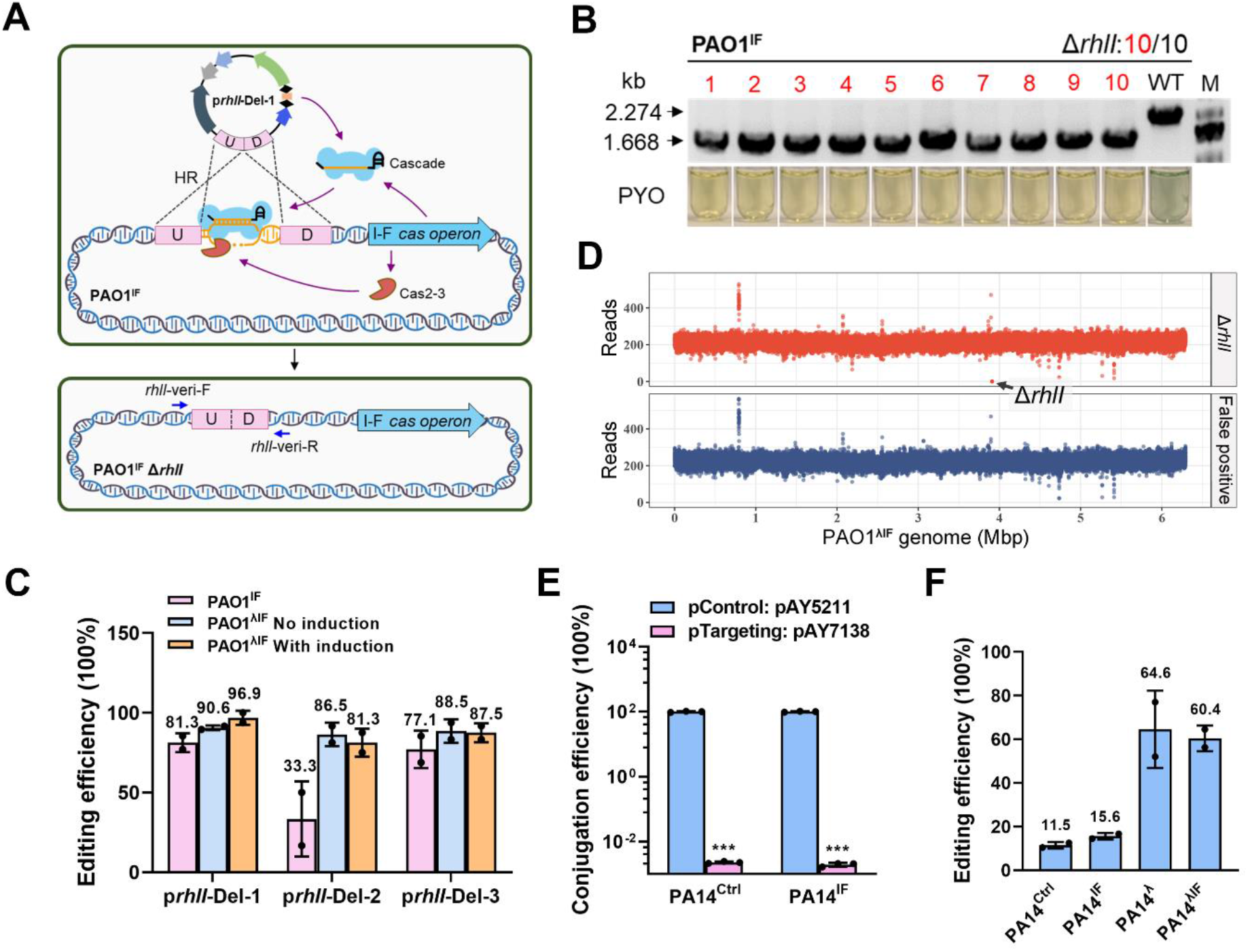
Transferred I-F Cascade-mediated precise and efficient gene deletion in PAO1 and PA14. (**A**) Schematic diagram of the transferred I-F Cascade-mediated genome editing by one-step transformation of a single editing plasmid p*rhlI*-Del-1. The editing plasmid expresses a *rhlI*-targeting crRNA and provides a donor template which consists of 833-bp upstream (U) and 805-bp downstream (D) homologous arms of the *rhlI* gene for homologous recombination (HR)-mediated repair. Cascade targeting recruits Cas2-3 to generate DNA breakage within the *rhlI* gene. HR between the donor and its homologous arms flanking the *rhlI* gene results in the desired *rhlI*-deletion mutant. (**B**) Verification of the precise *rhlI* deletion in the PAO1^IF^ host by colony PCR. Primer pairs used to verify *rhlI* deletion in this study are indicated in (a). Clone numbers with desired *rhlI* deletion are highlighted in red. Production (WT) or loss (mutant) of the blue-green colored pigment PYO in the corresponding inoculum is shown in the lower panel. M: Marker. (**C**) Editing efficiency of the transferred I-F (PAO1^IF^) or λ_red_-I-F (PAO1^λIF^) system employing the indicated editing plasmids (x-axis) with or without L-arabinose induction. (**D**) Representative mapped WGS reads of the authentic Δ*rhlI* mutant (red) and false positive clone (blue) against the PAO1^λIF^ genome. Arrow indicates the absence of reads at the *rhlI* locus in the Δ*rhlI* clone. (**E**) DNA interference activity of the native (PA14^Ctrl^) and transferred (PA14^IF^) I-F Cascades determined by the conjugation efficiency of transforming pAY7138 that targets to IF_Ps1 (Figure 2A). (**F**) Editing efficiency of the native and transferred I-F Cascade systems in the indicated host strains. PA14^Ctrl^: PA14 strain integrated with the empty transferrable system lacking the *cas* operon; PA14^IF^: PA14 strain integrated with the transferrable I-F Cascade; PA14^λ^: PA14 strain integrated with the transferrable system carrying the λ-Red recombination system; PA14^λIF^: PA14 strain integrated with the transferrable λ_red_-I-F Cascade. Data are the mean of two (C and H) or three (G) biological repeats and are expressed as mean ± S.D. Statistical significance is calculated based on student’s *t* test (***, *P* < 0.001).

We also examined the editing capacity of the transferred I-F Cascade mediated by a less effective crRNA, i.e. that targets to the protospacer sequence IF_Ps7 (conjugation efficiency of 0.0016% vs. 0.0002% for IF_Ps6). Delivery of the corresponding editing plasmid p*rhlI*-Del-2 into PAO1^IF^ by conjugation resulted in 33.3% editing efficiency (Figure 3C), indicating that the loci structure of the selected targeting site (protospacer) affects editing efficiency. Editing capacity of the transferred I-F Cascade by employing a tandem mini-CRISPR cassette (p*rhlI*-Del-3) which simultaneously targets to the IF_Ps6 and IF_Ps7 protospacers (Supplementary Figure S3A) was determined to be 77.1% (Figure 3C), suggesting simultaneously targeting to multiple protospacers represents an effective strategy to circumvent the inefficient targeting effected by one site.

We next examined the site specificity of the transferred I-F Cascade-mediated genome editing and the possible causes of false positive clones. Three clones which have been verified to harbor Δ*rhlI* and three false positive clones which displayed luminescence but was shown to contain an unedited *rhlI* allele were subjected to WGS analysis. Precise *rhlI* deletion without off-target mutation was verified in all three authentic Δ*rhlI* clones (Figure 3D), demonstrating that the type I-F Cascade-mediated genome editing is highly site-specific. Interestingly, in all three false positive clones, mutations in the *cas* genes were identified, i.e. 33-bp deletion in *cas2-3* in one clone and point mutations in *cas5* in two clones (Supplementary Table S2), which cause inactivation of the nuclease Cas2-3 and premature termination of Cas5, respectively. These results indicated that mutations in *cas* genes represent a major reason for the emergence of false positive clones.

To test the robustness of the transferred I-F Cascade to construct other types of genetic editing, gene (or DNA fragment) insertion such as N-terminal *FLAG-tagging* of *mexF* gene and C-terminal *gfp-tagging* of *rhlA* gene, and a C54T point mutation in *rhlI* were carried out (Supplementary Figure S4A and S4B). Randomly screening ten luminescence-positive clones recovered from the conjugation of corresponding editing plasmid into PAO^IF^ identified 7/10, 5/10, and 1/10 desired clones for N-terminal *FLAG*-tagging of *mexF, gfp*-tagging to the C-terminal of *rhlA*, and the C54T point mutation in *rhlI*, respectively (Supplementary Figure S4C and S4D), confirming the robustness of the transferred I-F Cascade for genetic manipulations.

### An advanced transferrable λ_red_-I-F system enables genome editing in genotypes with a poor intrinsic homologous recombination capacity

We next expand the application of the transferrable I-F Cascade to PA14, a strain with an existing native type I-F CRISPR-Cas system in its genome. We first compared the DNA interference activity of the native I-F Cascade in PA14 and that with an additional Cascade from PA154197 integrated at its *attB* site in the derivative PA14^IF^. Introduction of a self-targeting plasmid pAY7138 (targeting to the protospacer IF_Ps1) resulted in a conjugation efficiency of 6-magnitude lower relative to the control plasmid pAY5211 in both strains (Figure 3E), indicating the strong DNA interference of the native I-F system in PA14 and that presence of an additional I-F Cascade did not further enhance the DNA interference activity in the resulting cells. Surprisingly, when the editing plasmid p*rhlI*-Del-3 was delivered into the two cells, PA14 and PA14^IF^, to construct *rhlI* gene deletion, a very low editing efficiency (11.5% and 15.6%) was obtained in both cells (Figure 3F), implying that the recombination capacity might be poor in PA14.

We then transferred the advanced λ_red_-I-F cassette (pAY7136) we developed which contains both the PA154197 I-F Cascade and a λ-Red recombination system driven by P_BAD_ into PA14 and exploited the system to construct *rhlI* deletion. Upon induction of the λ-Red system by 20 mM L-arabinose, an editing efficiency of 60.4% was achieved for constructing *ΔrhlI* employing the editing plasmid p*rhlI*-Del-3 (Figure 3F), which is 4-fold higher than that achieved in the PA14^IF^ host (15.6%). This result demonstrated the advantage of the transferrable λ_red_-I-F cassette which enables the I-F Cascade-mediated genome editing in a recombination-poor genotype. The advantage of the transferrable λ_red_-I-F system was also demonstrated in the PAO1 background as evidenced by an increase of editing efficiency from 33.3% to more than 80% for constructing Δ*rhlI* in the PAO1^λIF^ host relative to the PAO1^IF^ host employing the same editing plasmid p*rhlI*-Del-2 (Figure 3C), suggesting that a greater homologous recombination capacity can improve the editing efficiency by compensating for the inefficient targeting of a programmed crRNA.

### The transferrable I-F Cascade is readily applied in diverse genetic backgrounds

To examine the widespread application of the transferrable λ_red_-I-F system, we integrated the λ_red_-I-F cassette into three clinical PA isolates with diverse genetic backgrounds and virulence capacities (Supplementary Table S3 and Supplementary Figure S5): PA150577, PA132533, PA151671, and characterized the DNA interference activity and editing exploitations of the system in the resulting cells. Since the genome sequence of PA150577 is available, we first conducted a CRISPRCasFinder (43) analysis and identified a complete native type I-F CRISPR-Cas loci in this strain which is organized in a sandwich structure of CRISPR1-Cas-CRISPR2 and shares 98.69% identity with that in PA154197 (Supplementary Figure S6A). However, the DNA interference activity of this native CRISPR-Cas system was found to be poor as evidenced by only 2-magnitude reduction of colony recovery upon introduction of the targeting plasmid pAY7138 relative to the control plasmid (Figure 4A), as opposed to 5 to 6-magnitude reduction necessary for to achieve genome editing as described above. PA132533 and PA151671 are clinical isolates without available genome information. Hence, we first conducted PCR analysis to identify the presence of native I-F CRISPR-Cas systems in these genomes. Presence of all six I-F Cascade genes was identified in PA151671 (Supplementary Figure S6B). However, introduction of the targeting plasmid pAY7138 did not reveal any DNA interference activity in the cell, indicating that the native type I-F system in PA151671 is inactive (Supplementary Figure S6C). In PA132533, PCR analysis indicated that the strain contains an incomplete I-F Cascade which lacks the *cas8f* gene (Supplementary Figure S6B).

**Figure 4.**
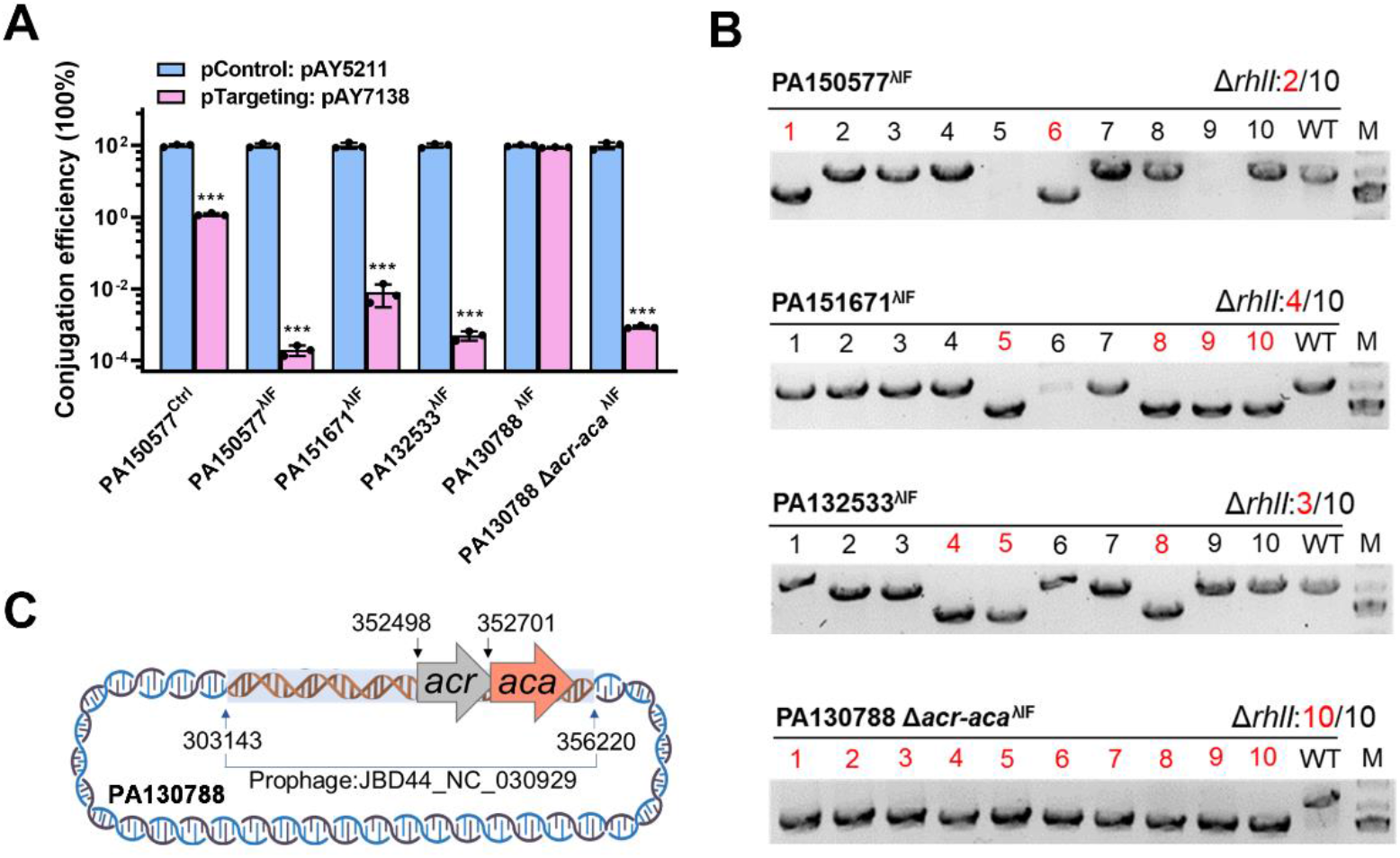
Transferrable λ**red**-I-F Cascade enables gene deletion in *P. aeruginosa* strains with diverse genetic backgrounds. (**A**) DNA interference activity of the transferred I-F Cascade determined by the conjugation efficiency of transforming pAY7138 (targeting to the IF_Ps1 protospacer site) into the indicated host strains PA150577^ctrl^, PA150577^λIF^, PA151671^λIF^, PA132533^λIF^, PA130788^λIF^, and PA130788 Δ*acr*-*aca*^λIF^ (x-axis). (**B**) Verification of the precise *rhlI* deletion in the indicated host strains PA150577^λIF^, PA151671^λIF^, PA132533^λIF^, and PA130788 Δ*acr*-*aca*^λIF^ by colony PCR. Primer pairs used to verify *rhlI* deletion in this study are indicated in Fig. 3a. Clone numbers with desired *rhlI* deletion are highlighted in red. (**C**) Schematic diagram of an anti-CRISPR gene *acr* and its associated repressor gene *aca* located in the prophage JBD44_NC_030929 region in PA130788. Data are the mean of three biological repeats and are expressed as mean ± S.D. Statistical significance is calculated based on student’s *t* test (***, *P* < 0.001).

We then transferred the λ_red_-I-F cassette into these three strains. Stable integration and expression of the cassette was confirmed by the blue colonies on the X-gal plate and RT-qPCR analysis (Supplementary Figure S7). Sufficiently high DNA interference activities were detected in all three resulting strains PA150577^λIF^, PA132533^λIF^ and PA151671^λIF^ (Figure 4A). Genome editing in the resulting cells was conducted by directly introducing the editing plasmid p*rhlI*-Del-3. A successful rate of 2/10, 4/10 and 3/10 was achieved in one-step, single conjugation reaction in the PA150577^λIF^, PA132533^λIF^ and PA151671^λIF^cells, respectively (Figure 4B). Together, these results demonstrated the powerful editing capacity and efficiency of the transferrable λ_red_-I-F Cascade we developed which effected editing in strains with and without native CRISPR-Cas systems, and strains with and without available genome sequences.

### Application of the transferrable and integrative I-F Cascade in *acr*-containing strains

Anti-CRISPR (Acr) is a common mechanism that inhibits the CRISPR-Cas immunity (44,45). More than 30% of sequenced *P. aeruginosa* genomes were found to carry one or more *acr* genes (20). We examined four additional *P. aeruginosa* strains, including two clinical isolates ATCC 27853 (termed as PA27853 herein) and PA130788 and two environmental strains PA238 and PA1155 in which presence of *acr* genes were indicated by the Acr prediction program AcrFinder (46) (Supplementary Figure S8). The λ_red_-I-F cassette was readily integrated into the *attB* site of the four genomes and were stably expressed as shown by RT-qPCR analysis (Supplementary Figure S7). However, no DNA interference activity was detected in all four strains, confirming the presence of Acrs. We next used PA130788 as a model to examine whether deleting *acrs* will activate the activity of the transferred Cascade. We deleted the *acr* gene together with its associated gene *aca* in PA130788 (Figure 4C) using the counter selection-based method (47), generating a mutant PA130788 Δ*acr*-*aca*. As shown in Figure 4A, the transferred λ_red_-I-F Cascade in this mutant was activated to execute self-targeting DNA breakage as evidenced by a 5-magnitude reduction of the colony recovery rate upon introduction of the pTargeting plasmid pAY7138 relative to the control plasmid pAY5211, confirming that the failure of self-targeting in PA130788^λIF^ was caused by the identified Acr. Construction of Δ*rhlI* by introducing the single editing plasmid p*rhlI*-Del-3 in PA130788 Δ*acr-aca* achieved 10/10 editing efficiency (Figure 4B). These results demonstrated the applicability of the transferred λ_red_-I-F system in *acr*-containing strains upon removal this element by an alternative approach.

### Repurposing the transferred I-F Cascade for transcriptional modulation

In addition to anti-CRISPR elements, inability to transfer plasmids due to poor DNA uptake capacity or the lack of suitable replicating vector in certain genotypes also impedes genome editing (11). Multidrug resistance in clinical strains poses another barrier for delivery of editing plasmids and selection of transformants. PA27853 represents such a genotype in which the CRISPR-Cas-based genome editing is hindered by both Acrs and its intrinsic resistance to kanamycin which restrained the transformation and selection of the series of targeting and editing plasmids we developed. To overcome these impediments, we set out to develop an ‘all-in-one’ transferrable I-F Cascade-based transcriptional interference system mini-CTX-CRISPRi-*lacZ* (transferrable CRISPRi) for transcriptional modulation without the necessary of delivering an editing plasmid. The transferrable I-F Cascade-based CRISPRi system (pAY6925) was constructed by removal of the *cas*2-3 gene in the transferrable I-F Cascade (pAY6924) to disable its DNA cleavage ability (Figure 5A). It is expected that with the provision of a programmed crRNA, the Cascade complex is able to recognize and bind to the complement genomic locus specifically, resulting in transcriptional repression by preventing the recruitment of RNA polymerase (RNAP) or blocking the transcription of the target gene (Figure 5B) (48). To bypass the delivery of additional plasmids such as that expressing a programmed crRNA for desired genomic site targeting, we designed a multi-cloning site downstream of the P*tat* promoter in the mini-CTX-CRISPRi-*lacZ* backbone such that a mini-CRISPR can be readily assembled into the integrative cassette, and upon integration of the entire cassette into the strains of interest, the crRNA will be constitutively expressed from the P*tat* promoter, eliciting robust Cascade binding to the target site (Figure 5A).

**Figure 5.**
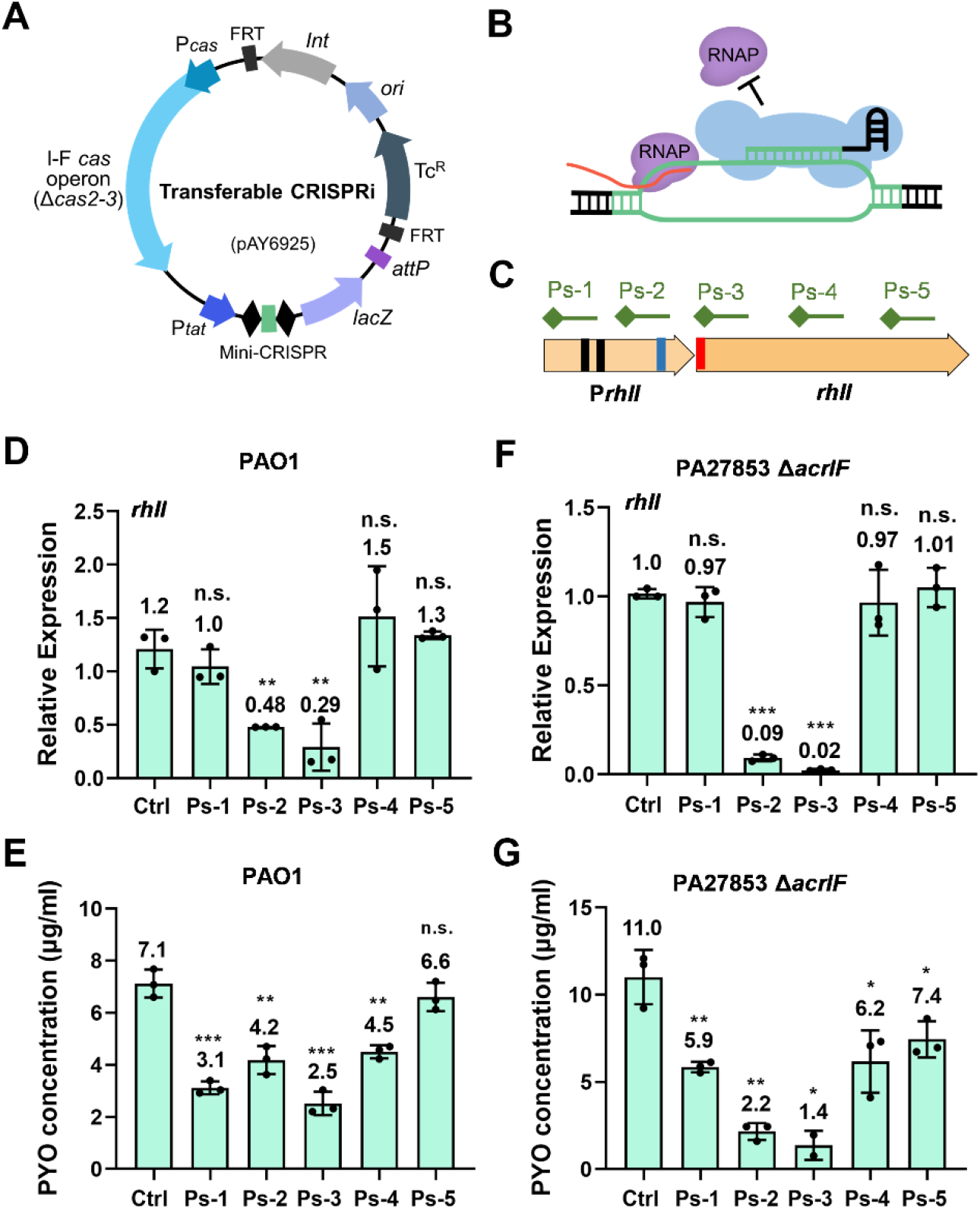
Transferrable I-F CRISPRi-mediated gene repression. (**A**) Diagram for the contents and organization of the transferrable I-F CRISPRi system. A truncated *cas* operon lacking the *cas2-3* gene replaced the complete *cas* operon as in the transferrable I-F Cascade (pAY6924). A programmed mini-CRISPR is assembled downstream of the P*tat* promoter. (**B**) Schematic diagram showing the working mechanism of the CRISPRi-based gene repression. Site-specific binding of the Cascade to a genomic locus prevents the recruitment of the RNA polymerase (RNAP) or blocks its operation in gene transcription. (**C**) Schematic diagram of the protospacers selected to examine the gene repression activity of the transferred CRISPRi system. Five protospacers located at different locations in the coding region of *rhlI* and its promoter region are selected. Diamonds denote the PAM sequences. Ps-1: −152 ~ −120 bp; Ps-2: −63 ~ −31 bp; Ps-3: 54 ~ 86 bp; Ps-4: 238 ~ 270 bp; Ps-5: 520 ~ 552 bp. (**D, E**) Relative expression of *rhlI* (D) and PYO production level (E) in PAO1 cells transferred with a I-F CRISPRi targeting to the indicated sites (x-axis). (**F, G**) Relative expression of *rhlI* (F) and PYO production level (G) in PA27853 *ΔacrIF* cells transferred with a I-F CRISPRi targeting to the indicated sites (x-axis). Data are the mean of three biological repeats and are expressed as mean ± S.D. Statistical significance is calculated based on student’s *t* test (n.s., Not significant; *, *P* < 0.05; **, *P* < 0.01; ***, *p* < 0.001).

We first tested the applicability of the transferrable I-F Cascade-based CRISPRi system in the model strain PAO1. We designed five transferrable mini-CTX-CRISPRi-*rhlI* plasmids, each expressing a crRNA targeting to a distinctive location in the *rhlI* gene and its promoter region (Figure 5C), including a region bound by the RNAP (Ps-1, −152 ~ −120 bp), transcription initiation region (Ps-2, −63 ~ −31 bp), the 5’-end of the *rhlI* coding region (Ps-3, 54 ~ 86 bp), the middle region of the *rhlI* gene (Ps-4, 238 ~ 270 bp), and the 3’-end of the *rhlI* coding region (Ps-5, 520 ~ 552 bp). We assembled each of the five mini-CRISPRs into the transferrable I-F Cascade CRISPRi backbone, respectively, resulting in five CRISPRi vectors, mini-CTX-CRISPRi-*rhlI*-1 to 5. Introduction of these vectors resulted in simultaneous integration of the I-F CRISPRi Cascade and a fragment expressing a *rhlI*-targeting crRNA at the *attB* site of PAO1. Three clones of each of the resulting constructs were selected to quantify the *rhlI* transcriptional levels. Among them, the construct integrated with the mini-CRISPR-2 and mini-CRISPR-3 displayed significant (51% and 71% reduction, respectively) repression of *rhlI* expression (Figure 5D). crRNAs expressed from these two mini-CRISPRs target to the regions located at the transcription initiation site (Ps-2) and the 5’-end of the *rhlI* coding region (Ps-3). PYO levels in these cells are consistent with the corresponding *rhlI* gene repression (Figure 5E). No repression effect was observed by targeting to Ps-1, Ps-4 and Ps-5. Interestingly, although RT-qPCR showed no significant change on *rhlI* expression by targeting to Ps-1 and Ps-4, PYO level was reduced moderately in the cell, suggesting certain degree of repression. These results demonstrated the feasibility of the designed CRISPRi system for gene repression by one-step transfer and integration of the I-F CRISPRi Cascade together with a programmed mini-CRISPR. It also suggested that most effective repression requires the Cascade targeting at the transcription initiation site or its proximal region.

We next tested the applicability of the system in PA27853. To this end, the *acr* gene in PA27853 was firstly deleted using the two-step counter-selection method (47). Each of the five transferrable mini-CTX-CRISPRi-*rhlI* plasmids was delivered into PA27853 *ΔacrIF* by conjugation. RT-qPCR and PYO production verified significant repression of *rhlI* effected by the cassette containing mini-CRISPR-2 and mini-CRISPR-3 (Figure 5F and 5G). This result demonstrated that the transferrable and integrative I-F CRISPRi system we developed provides as a powerful and an alternative gene knockdown strategy for functional genomics without the need of plasmid transformation. The robustness of this system was further demonstrated in the strain PA154197 Δ*cas2*-*3* and an additional clinical isolate PA153301 which genome sequence is not available but was verified to be free of a complete endogenous type I-F CRISPR-Cas system, i.e. only the presence of *cas6* gene was detected (Supplementary Figure S6B). In both strains, one-step transformation and integration of the mini-CTX-CRISPRi-*rhlI* containing mini-CRISPR-2 and −3 resulted in repression of *rhlI* (Supplementary Figure S9).

### Removal of the transferred λ_red_-I-F cassette

To generate clean gene deletion in the genetic background of interest, it is necessary to remove the transferred I-F Cascade. Although the mini-CTX-*lacZ* backbone was engineered with two FRT sites for the removal of the auxiliary elements in the vector such as the integrase gene *Int* and the tetracycline-resistant marker employing the ectopically expressed flippase (39), the I-F Cascade and the *lacZ* reporter remains at the *attB* site. To completely remove the integrated fragment following the desired genetic exploitations, we harnessed the processive degradation capacity of the Cas3 nuclease in the transferred I-F Cascade to delete the large-scale fragment of λ_red_-I-F Cascade (21.212 kb). The *lacZ* reporter included in the fragment serves as an indicator for successful removal of the whole cassette with a desired white colony phenotype on the X-gal plate. A 22.298-kb Cascade-removal plasmid pAY7401 (Figure 6A) was constructed by assembling a crRNA targeting to the *lacZ* gene and a donor sequence consisting of ~5-kb upstream and ~5-kb downstream homologous arms of the *attB* site. We tested the capacity of this editing plasmid in the PAO1^λIF^ strain by introducing pAY7401 *via* conjugation and selected white colonies on the X-gal plates. As shown in the Supplementary Figure S4E, selective white colonies were recovered, suggesting the successful removal of the integrated *lacZ*-containing λ_red_-I-F cassette. Successful removal of the integrated fragment was also confirmed by PCR amplification of the region flanking the *attB* site using primer pairs P1 and P2 located upstream and downstream of the donor sequences, respectively (Figure 6B), resulting in a fragment with the desired site of 10.3 kb (Supplementary Figure S4F). The capacity of the common cassette-deleting editing plasmid was also verified by introducing pAY7401 into PAO1^λIF^ Δ*rhlI* cell to delete the entire mini-CTX-λ_red_-IF-P*tat-lacZ* following curing the editing plasmid p*rhlI*-Del-1 (Supplementary Figure S4G). Colony PCR verified the successful removal of the cassette by the size of the PCR product (Figure S6B). The resulting clones were also shown to be susceptible to tetracycline, the antibiotic resistant marker in the mini-CTX-*lacZ* backbone (Supplementary Figure S4E). Meanwhile, the Δ*rhlI* genetic allele remains unchanged during the process as verified by PCR analysis. Together, these results demonstrated that the transferrable λ_red_-I-F Cascade is capable of not only effecting diverse genetic editing in the heterologous hosts, but also being exploited to remove large-scale genomic fragments (>20 kb) with convenient selection.

**Figure 6.**
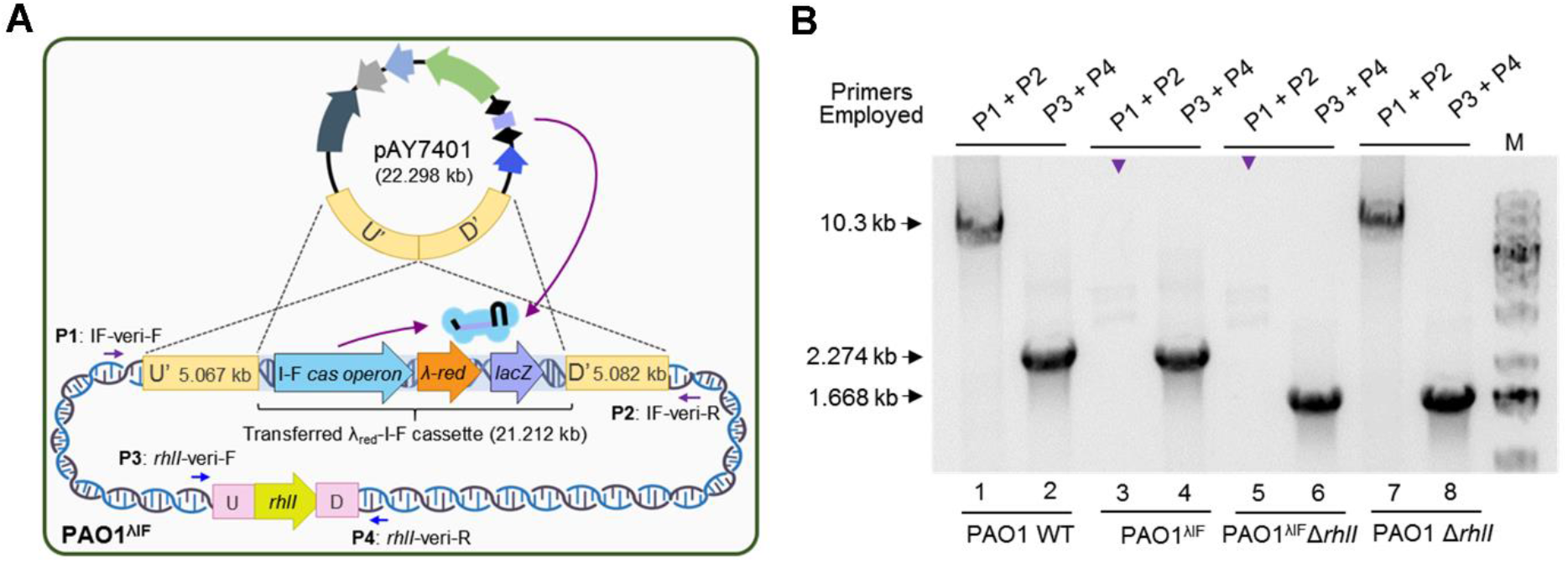
Removal of the transferred λ_red_-I-F cassette. (**A**) Schematic diagram showing the working mechanism of deleting the transferred λ_red_-I-F cassette by one-step transformation of a single editing plasmid pAY7401. The editing plasmid expresses a *lacZ*-targeting crRNA and provides a donor template which consists of 5.067-kb upstream (U’) and 5.082-kb downstream (D’) homologous arms of the *attB* site for homologous recombination. (**B**) PCR analysis to verify the desired gene editing and removal of the transferred λ_red_-I-F cassette. Primer pairs P1 and P2 are used to verify removal of the transferred λ_red_-I-F cassette. Primer pairs P3 and P4 are used to verify *rhlI* deletion. Presence of a 10.3-kb PCR product using the P1 and P2 primer pair indicated absence of cassette integration at the *attB* site, i.e., in the PAO1 WT and the clean edited cell PAO1 *ΔrhlI*. Absence of a PCR product using P1 and P2 primer pair indicates the presence of the integrated cassette which was incapable to be amplified by our PCR polymerase due to its large size.

## DISCUSSION

Despite that class 1 CRISPR-Cas systems represent 90% of all naturally occurring CRISPR-Cas systems, their exploitations for heterologous genome editing have been hindered by the complexity of the effector Cascade. In the present study, we established a transferrable I-F Cascade based on the integration-proficient mini-CTX vector which enables integration of exogenous DNA fragments at the conserved *attB* site in diverse *P. aeruginosa* genomes for stable expression and function, converting CRISPR-free cells or those containing an inactive or degenerated CRISPR-Cas system into cells with an endogenously active type I-F Cascade. Exploiting this stably expressed ‘native’ type I-F Cascade, efficient and precise genome editing was achieved by introducing a single editing plasmid in one step. An advanced transferrable λ_red_-I-F Cascade is further developed to simultaneously integrate an I-F Cascade and a λ-Red recombination system in the heterologous hosts, improving genome-editing efficiency in recipient cells with a poor intrinsic HR capacity. The transferred λ_red_-I-F Cascade can also be harnessed for deleting large-scale genomic fragments (>20 kb) such as the entire transferred cassette, resulting in clean genetic manipulations in the genetic background of interest. Applications of the system are demonstrated in diverse genetic backgrounds, including strains with poor HR capacities, with or without native CRISPR-Cas systems, wild clinical isolates without genome sequences, as well as strains containing Acrs. To our knowledge, this is the first transferrable and integrative type I Cascade for widespread and robust heterologous genome editing. Lastly, a reprogrammable ‘all-in-one’ I-F Cascade-based CRISPRi system was developed to provide a powerful and alternative gene knockdown strategy for functional genomics without the need of delivering additional plasmids to express crRNA guides, further expanding the applicability of the transferrable system to a broad range of hosts with poor DNA uptake capacity and antibiotic-resistance. The methodology provides a framework to expand the diverse type I Cascades for widespread, heterologous genome editing.

The transferred I-F Cascade displayed strong DNA interference activity and efficient editing capacity in diverse genetic backgrounds. Nonetheless, false positive clones emerge following the editing reactions. Our WGS analysis showed that all three randomly selected false positive clones recovered from the editing reaction to construct *ΔrhlI* carry mutations in the *cas* genes, i.e. *cas5* and *cas2-3*. In another attempt to construct *ΔpqsA* in PAO1^λIF^ (data not shown), we identified false positive clones carrying mutated editing plasmid in which the spacer and the downstream repeat sequence was excised (Supplementary Figure S10), suggesting mutations in *cas* genes or the self-targeting mini-CRISPR in the editing plasmid are the major reasons for the emergence of false positive clones (49). Hence, future efforts to increase the editing efficiencies should focus on preventing these spontaneous mutations, such as modifying the repeat sequence in the mini-CRISPR to prevent the spacer excision *via* homologous recombination between the two repeats (15).

Our studies also indicated that the loci structures of protospacers being targeted affect the editing efficiency. For instance, an editing efficiency of 81.3% and 33.3% was achieved when pEditing expressing a crRNA targeting to the IF_Ps6 and IF_Ps7 site, respectively, was employed for constructing Δ*rhlI*. These results suggest that comprehensively investigating the sequence preference of a Cascade may facilitate the selection of the most effective protospacer for genome editing. Nonetheless, we demonstrated that simultaneously targeting to multiple protospacers using a tandem mini-CRISPR is effective to prevent the inefficient targeting and editing by targeting to only one locus. Moreover, we showed that increasing the HR capacity by supplying an exogenous recombination system improved the editing efficiency in the case a relatively inefficient targeting site was employed. This capacity will be highly desirable in certain editing applications which involve limited protospacer sequence and location selections, such as site-specific gene insertion or point mutation.

The widely distributed anti-CRISPR elements (Acrs) are regarded as another major obstacle limiting the CRISPR-Cas-mediated genome editing, especially in the wild clinical and environmental *P. aeruginosa* isolates (20). We identified Acrs in all four wild isolates (two clinical and two environmental strains) which showed no DNA interference activity following integration of the I-F Cascade. Activities of the transferred I-F Cascade in two strains PA130788 and PA27853 were activated upon deletion of the *acr* genes and highly efficient gene deletion and robust gene repression was achieved subsequently. Thus, by combining the counter selection-based gene-deletion method to remove *acr* genes, the application of the transferred I-F Cascade can be expanded in strains containing Acrs. Although removal of *acr* genes is relatively laborious and time-consuming using conventional methods, the method is still advantageous as the resulting cells will enable one-step transferrable Cascade-based genome editing or gene repression. An alternative strategy to overcome Acrs is to overexpress the anti-CRISPR-associated gene (*aca*) which represses *acr* and in turn actives CRISPR-Cas systems (50). This strategy has been demonstrated to be effective in a type I-C Cascade-mediated genome editing in a PAO1 variant which is lysogenized by a recombinant DMS3m phage expressing the anti-CRISPR gene *acrIC1* (15). However, robustness of Aca-mediated anti-anti-CRISPR strategy might be compromised in clinical strains owing to their complicated genetic backgrounds. We have applied this approach in PA130788 by overexpressing the *aca* gene of the identified *acr*. Despite that expression of *acr* was significantly repressed upon introduction of an ectopic *aca* gene and DNA interference activity of the transferred I-F Cascade was improved as shown by a conjugation efficiency of 0.5% upon introduction of pAY7138 (Supplementary Figure S11), we failed to obtain a desired Δ*rhlI* mutant after screening 960 recovered clones. Since the *acr* genes present in *P. aeruginosa* predominantly inhibit the three prevalent CRISPR-Cas subtypes I-C, I-E, and I-F in this species (45), it is conceivable that developing transferrable I-A or I-B Cascade-mediated genome editing approach may overcome the barrier imposed by the common Acrs in *P. aeruginosa*.

The type II Cas9 effector has been broadly employed for genome editing in various eukaryotic and prokaryotic hosts. However, the transferrable and integrative Cas9 system failed to effect efficient self-targeting in the various recipient hosts we tested in this study. In *P. aeruginosa*, a two-plasmid-based Cas9 approach was developed to achieve efficient genome editing in PAO1 and PAK (23). By modifying the editing plasmid in the two-plasmid system to contain an origin of transfer *(oriT)*, we improved the delivery efficiency of this plasmid by conjugation and successfully delivered the plasmid into PA14 and clinical strains PA151671 and PA132533. However, we found that the targeting efficiency mediated by the plasmid-borne Cas9 was still substantially lower than the transferred I-F Cascade in these strains especially in the clinical strains. This is probably due to the instability and degradation of the Cas9 protein in these cells (51). The success and ease of the transferred I-F Cascade-mediated genome editing in the clinical isolates PA151671 and PA132533 in which neither the transferred nor the plasmid-borne Cas9 effects any DNA interference clearly demonstrated the greater capacity and robustness of the transferred I-F Cascade than the Cas9 effector for heterologous genetic manipulations. Together, our studies demonstrated the power and potentials of transferrable type I Cascades for heterologous genome editing.

## DATA AVAILABILITY

All genome data are deposited in the NCBI under the project accession number PRJNA694101 and the transcriptome sequencing result is available on FigShare at https://doi.org/10.6084/m9.figshare.13699492

## SUPPLEMENTARY DATA

Supplementary Data are available at NAR Online.

## ACKNOWLEDGEMENTS

We thank Dr. Xin Deng (Department of Biomedical Sciences, City University of Hong Kong) for providing the mini-CTX-*lacZ* vector and thank Ms. Xiaojue Wang (School of Biological Sciences, The University of Hong Kong) for measurement of *cas* gene expression in the clinical strains.

## Supplementary data

**Figure S1.**
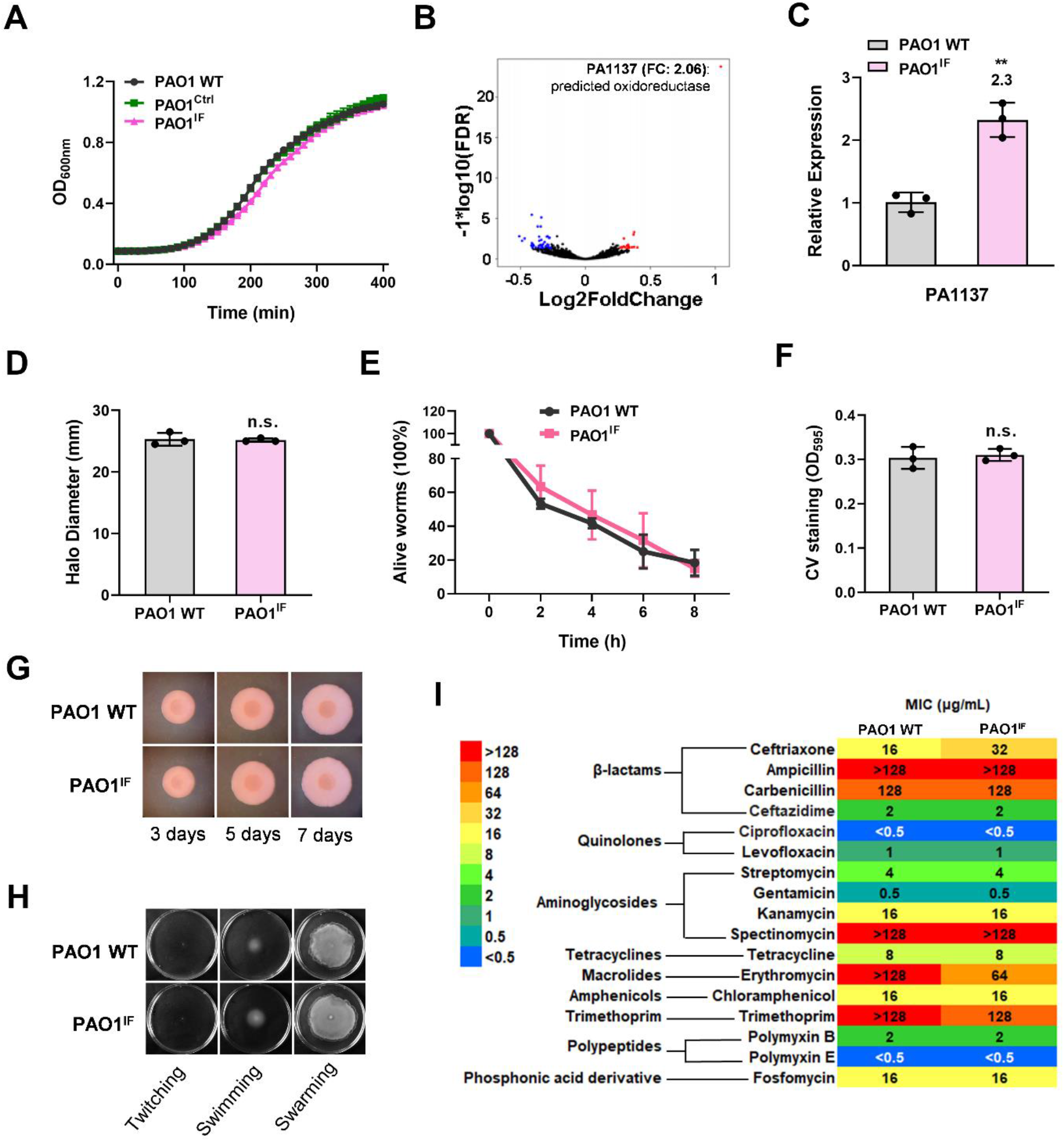
Integration of the transferrable I-F Cascade (mini-CTX-IF-P*tat-lacZ*) does not perturb the physiology of the recipient strain. Growth curve (**A**), transcriptome (**B**), exo-proteolytic activity (**D**), virulence (assessed by the *C. elegans* killing assay) (**E**), biofilm formation (**F**), colony biofilm morphology (**G**), motility (**H**) and antibiotic susceptibility (**I**) of the PAO1 wild-type (PAO1 WT) and PAO1^IF^ strains. (**C**) RT-PCR analysis of the expression of PA1137 which was shown to be ~2-fold higher in the PAO1^IF^ strain than in PAO1 WT by RNA-seq (**B**). Data are the mean of three biological repeats and are expressed as mean ± S.D. Statistical significance is calculated based on student’s *t* test (n.s., Not significant; **, *P* < 0.01).

**Figure S2.**
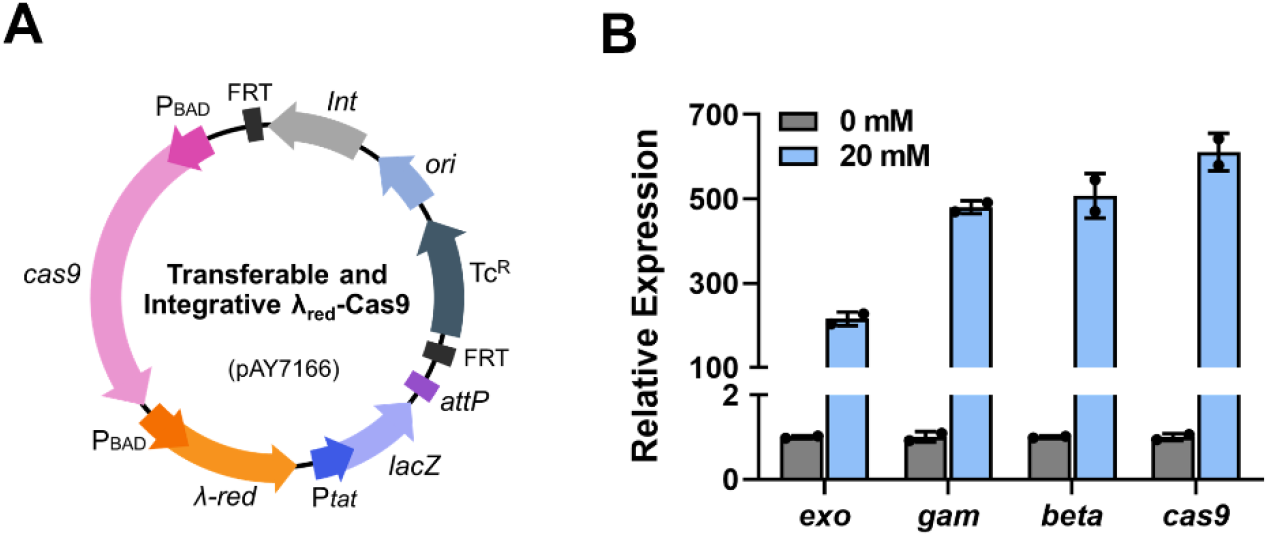
Construction and gene expression verification of the transferrable Cas9 system. (**A**) Diagram of the transferrable λ-Cas9 system. The system carries a *cas9* gene driven by the P_BAD_ promoter. (**B**) RT-qPCR showing the expression of the *cas9* and λ-Red genes in PAO1^λCas9^ upon induction by 20 mM L-arabinose.

**Figure S3.**
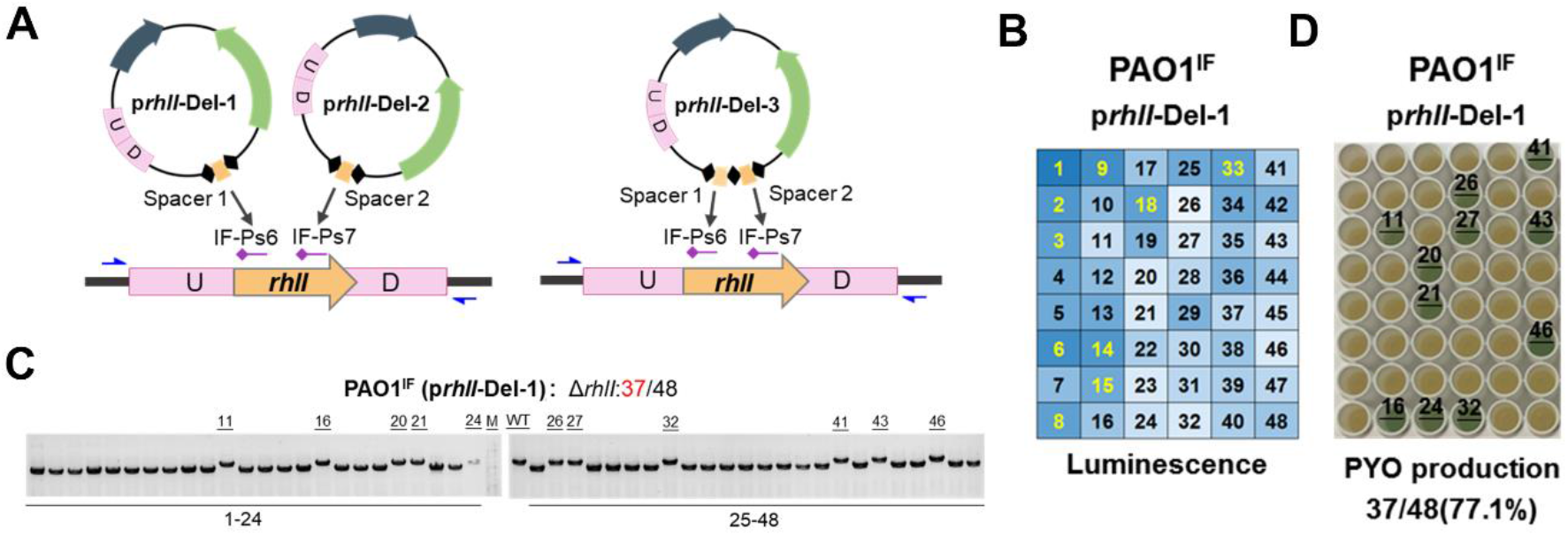
Construction of the *rhlI* deletion mutant by exploiting the transferred I-F Cascade in PAO1^IF^ and three editing plasmids. (**A**) Diagrams showing the organization and targeting sites of three editing plasmids employed for constructing *rhlI* deletion in PAO1^IF^. p*rhlI*-Del-3 contains two spacers simultaneously targeting to the IF-Ps6 and IF-Ps7 protospacer sites in the *rhlI* gene. (**B**) A representative panel of luminescence-based selection of clones recovered from delivery of the editing plasmid p*rhlI*-Del-1 into PAO1^IF^. 48 clones are inoculated. 10 clones displaying the highest luminescence intensity are indicated in yellow. (**C**) Colony PCR to verify *rhlI* deletion in 48 randomly selected clones recovered from the introduction of p*rhlI*-Del-1 into PAO1^IF^ (b). 37 out of 48 clones show desired size of the bands. 11 false positive clones are indicated. (**D**) A representative of PYO production screening to verify *rhlI* deletion in PAO1^IF^. Cell cultures of the corresponding 48 clones in (B) are shown. 37 positive clones display loss of blue-green colored PYO. 11 false positive clones were indicated in which PYO production (blue-green color) was unaffected.

**Figure S4.**
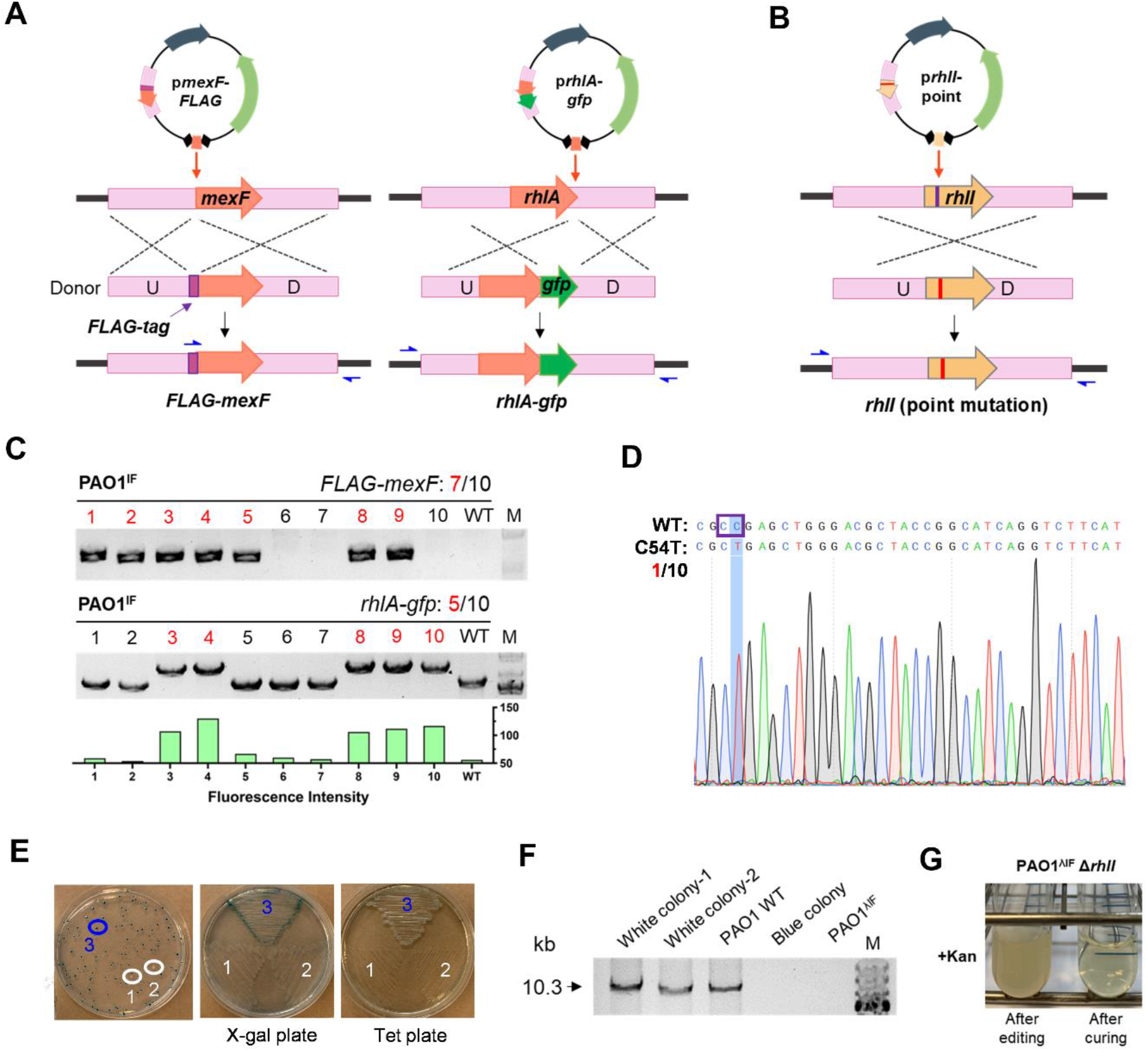
Harnessing the transferrable type I-F Cascade to achieve various genetic manipulations in PAO1^IF^ and PAO1^λIF^. (**A**) Schematic diagram showing the design of editing plasmids for N-terminal *FLAG*-tagging in *mexF (pmexF-FLAG)* and C-terminal *gfp*-tagging in *rhlA (prhlA-gfp*). For N-terminal *FLAG*-tagging (left panel): A 32-bp sequence spanning the start codon (ATG) of *mexF* is selected as the protospacer and the donor contains a 24-bp *FLAG* sequence immediately following the start codon. For C-terminal *gfp*-tagging (right panel): A 32-bp sequence spanning the stop codon (TGA) of *rhlA* is selected as the protospacer and the donor contains a 714-bp *gfp*-encoding sequence placed immediately upstream of the stop codon. Blue arrows indicate the primers used for verification. (**B**) Schematic diagram showing the design of editing plasmid (p*rhlI*-point) for a point mutation (C54T) in *rhlI*. A PAM sequence “5’-<u>C</u>53<u>C</u>54-3’” is selected to conduct C54T substitution. The donor sequence consists of 500-bp sequence which spans the target site and contains the desired C54T substitution. (**C**) Verification of the precise *FLAG*-tagging of *mexF* and *gfp*-tagging of *rhlA* in PAO1^IF^ host by colony PCR. Primer pairs used for PCR verification are indicated in (A). Clone numbers with desired *FLAG or gfp*-tagging are highlighted in red. Green fluorescence is detected in the clones with desired *gfp*-tagging. M: Marker. (**D**) A representative DNA sequencing result demonstrating the success of C54T point mutation in *rhlI*. The PAM sequence is framed by purple box. (**E**) A representative plate showing the recovery and selection for clones in which the transferrable system are removed (white clones). Two white clones were selected and streaked onto X-gal (40 μg/ml) and tetracycline (Tet, 50 μg/ml) −containing plates for further verification. (**F**) PCR analysis of the fragments flanking the *attB* site in various strains to verify the removal of the transferred λ_red_-I-F cassette (10.3 kb in PAO1 WT). (**G**) Curing of the editing plasmid in the edited cells. Edited cells could not grow in the presence of kanamycin (100 μg/ml) after plasmid curing.

**Figure S5.**
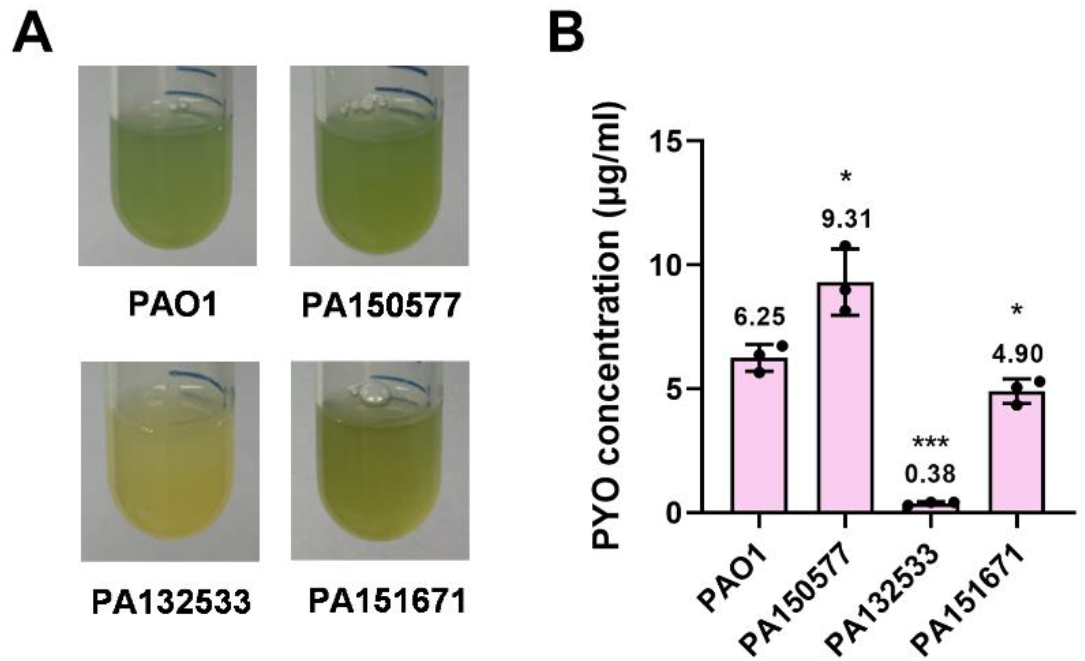
PYO production capacity of the clinical isolates PA150577, PA132533 and PA151671 (PAO1 serves as the reference strain). Production of the blue-green colored pigment PYO in the corresponding inoculum is shown in (A) and quantified in (B). Data are the mean of three biological repeats and are expressed as mean ± S.D. Statistical significance is calculated based on student’s *t* test (*, *P* < 0.05; ***, *P* < 0.001).

**Figure S6.**
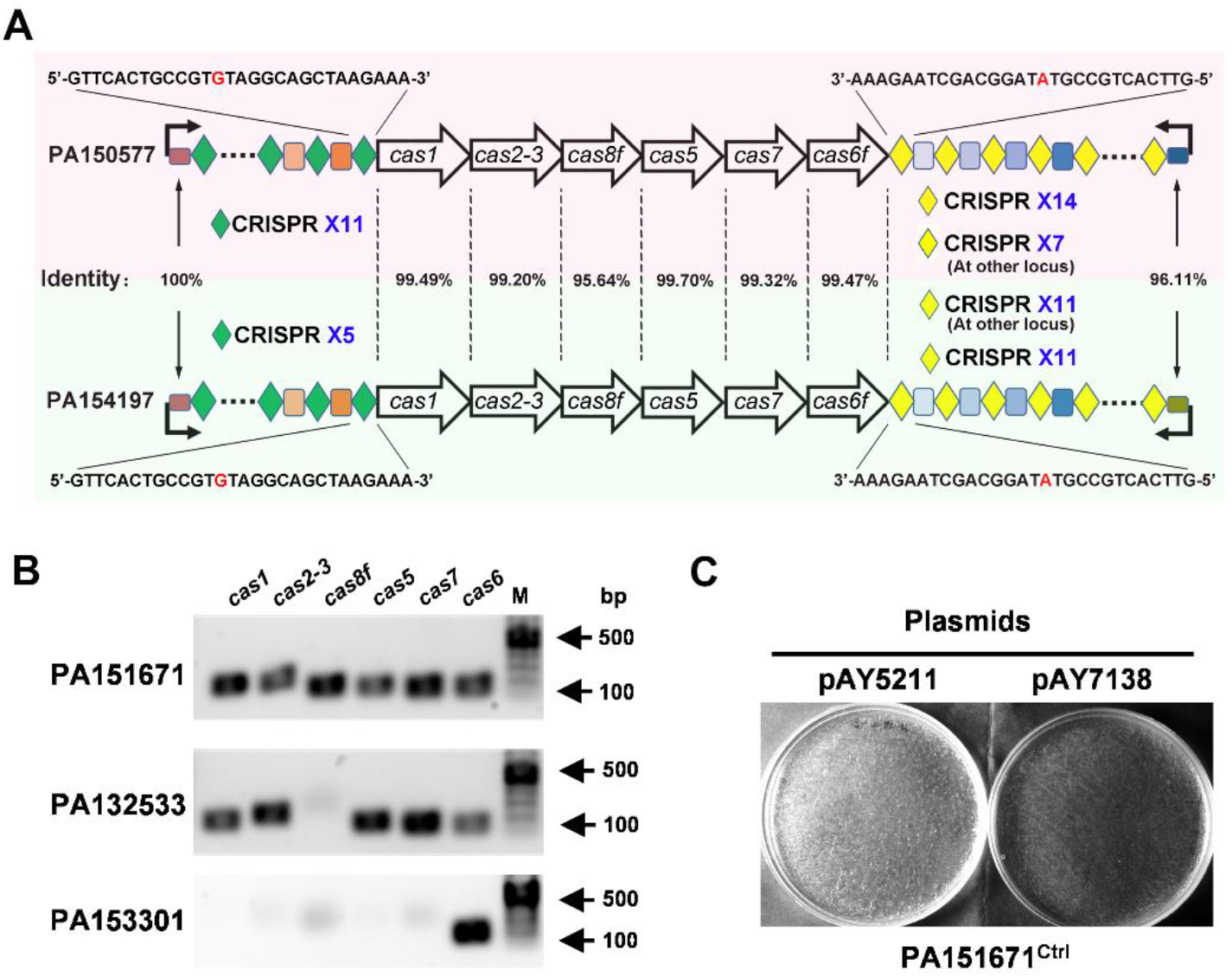
Characterization of the native I-F CRISPR-Cas systems in PA150577, PA151671, PA132533 and PA153301. (**A**) Comparison of the type I-F CRISPR-Cas system in PA150577 with that in PA154197. (**B**) PCR-based detection for the presence of I-F *cas* genes in the clinical isolates PA151671, PA132533 and PA153301. Primers used are the same as that used for RT-qPCR in Fig. 1i. PA132533 and PA153301 contain an incomplete type I-F CRISPR-Cas system in their genomes. (**C**) DNA interference activity of the native type I-F CRISPR-Cas system in PA151671 assessed by colony recovery following introduction of the targeting plasmid pAY7138. No obvious DNA interference was observed.

**Figure S7.**
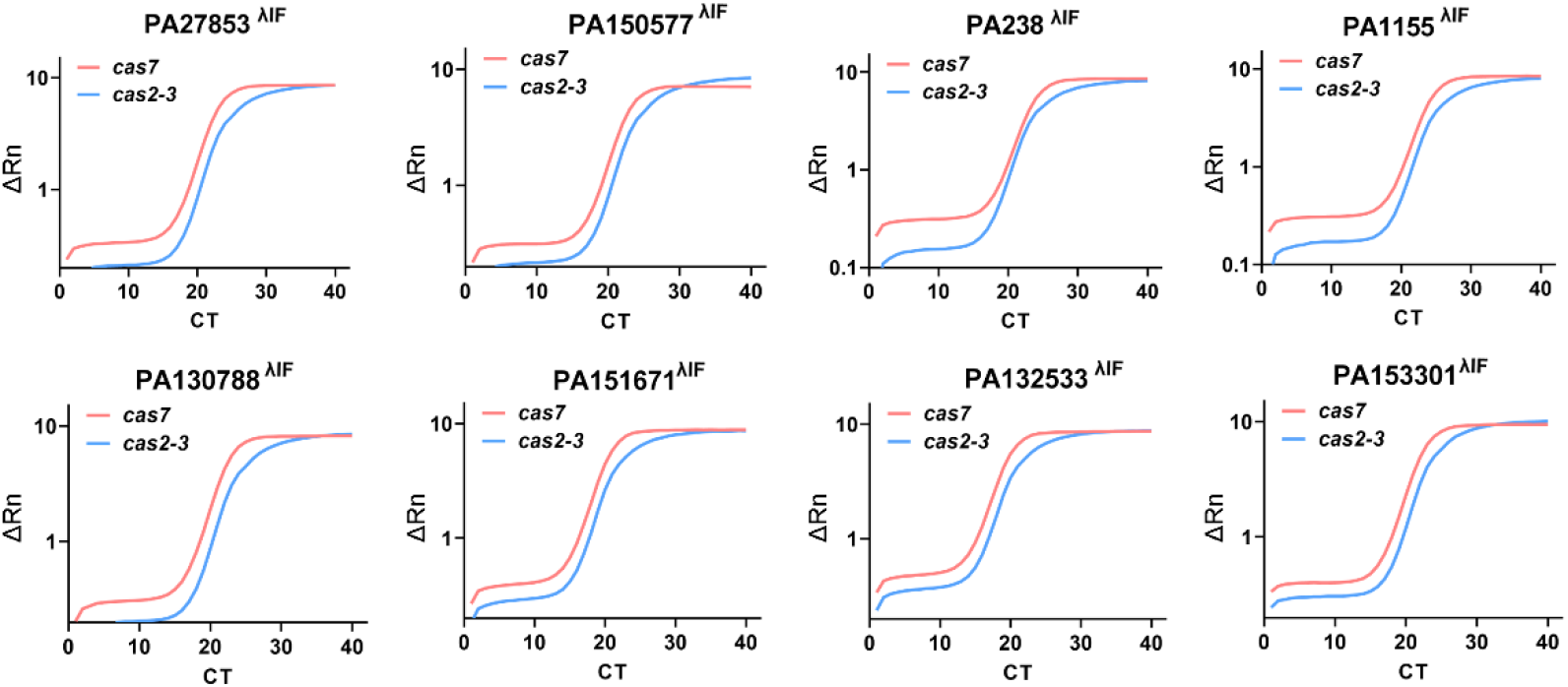
Integration and expression of the transferrable λ_red_-I-F Cascade in diverse *P. aeruginosa* recipient strains. Quantitative PCR showing the stable expression of the *cas2-3* and *cas7* genes in all the eight wild *P. aeruginosa* isolates transferred with λ_red_-I-F Cascade.

**Figure S8.**
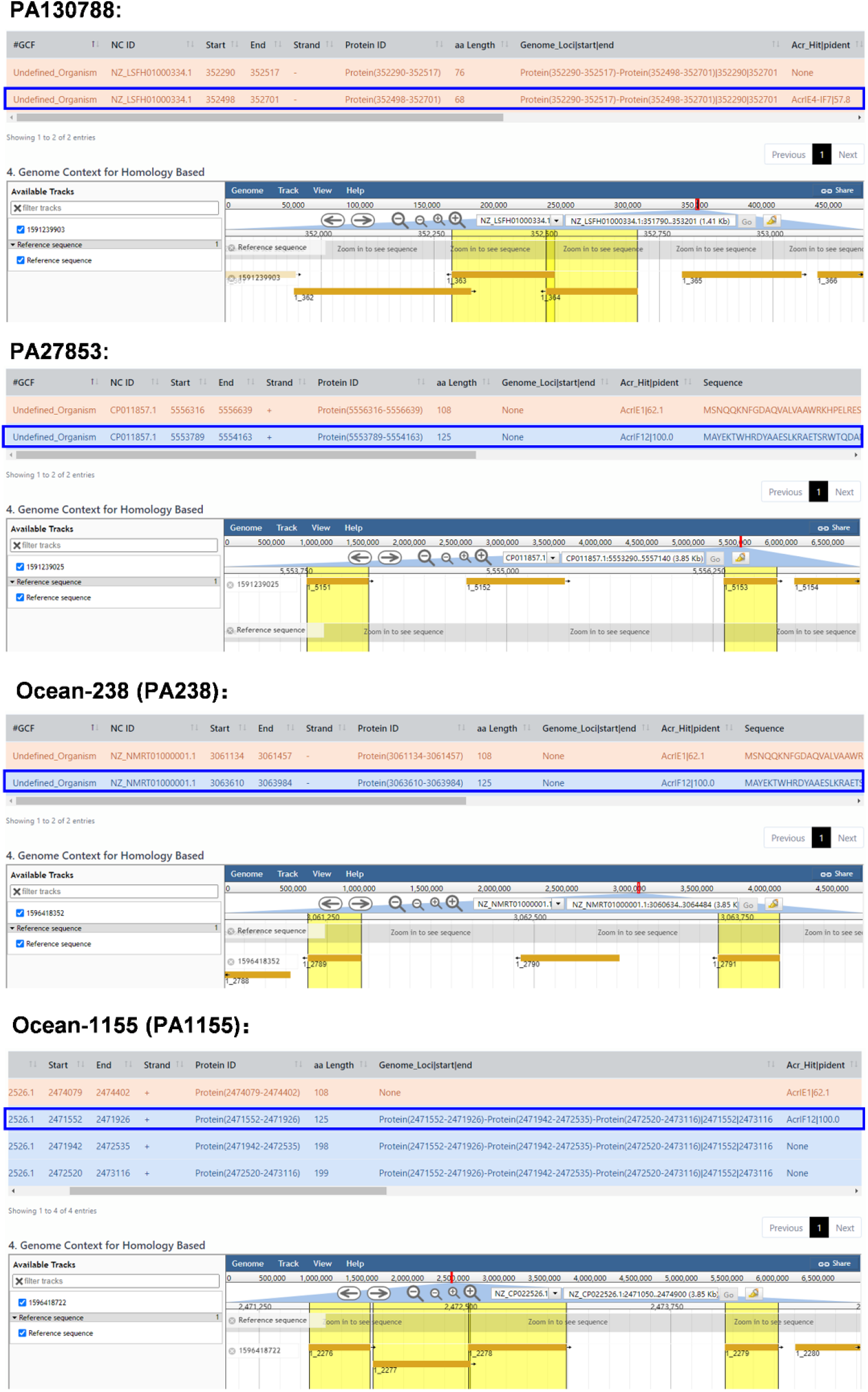
Bioinformatic analysis reveals the presence of anti-CRISPR gene (that potentially inactivates the type I-F CRISPR-Cas system) in the genome of PA130788, PA27853, PA238 and PA1155. Predicted *acr* genes in the host genomes are highlighted in yellow boxes (Lower panel of each strain). Acrs which have been demonstrated to inactivate the type I-F CRISPR-Cas system are framed in blue boxes (Upper panel of each strain).

**Figure S9.**
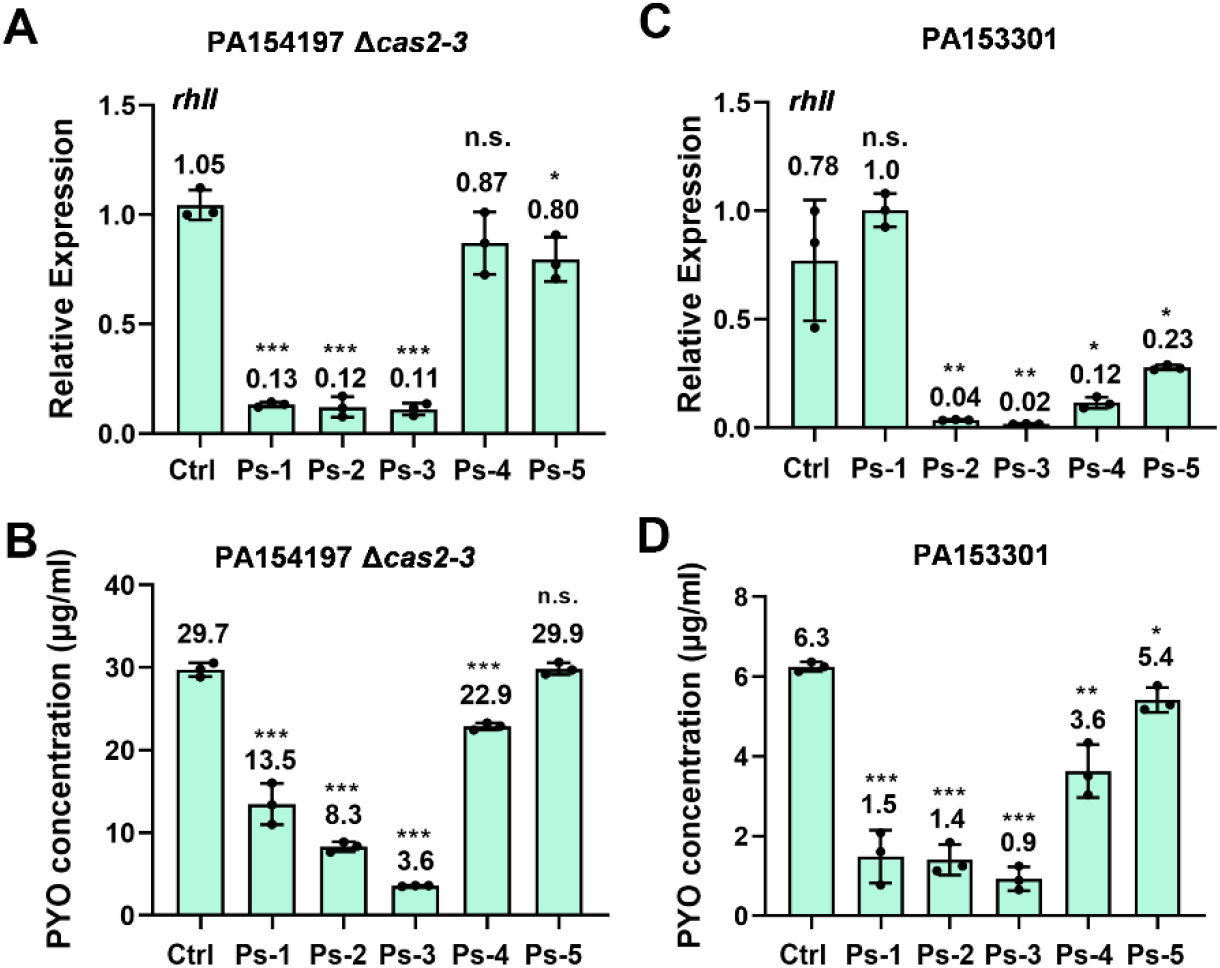
Transferrable I-F CRISPRi-mediated gene repression in PA154197 *Δcas2-3* and PA153301. (**A, B**) Relative expression of *rhlI* (A) and PYO production level (B) in PA154197 *Δcas2-3* cells transferred with a I-F CRISPRi targeting to the indicated sites (x-axis). (**C, D**) Relative expression of *rhlI* (C) and PYO production level (D) in PA153301 cells transferred with a I-F CRISPRi targeting to the indicated sites (x-axis).

**Figure S10.**
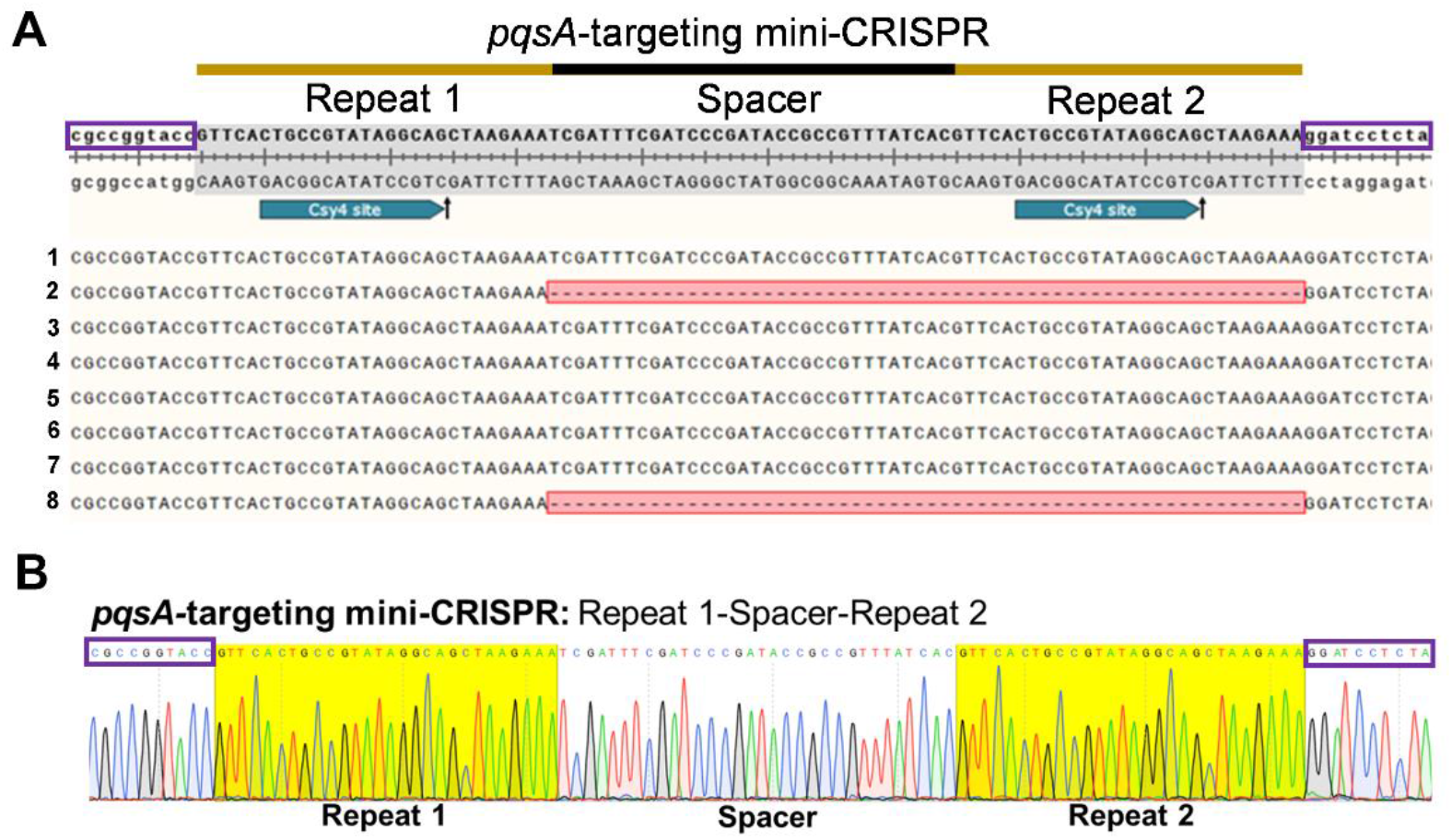
Mutations in the mini-CRISPR sequence targeting to the *pqsA* gene were identified in false positive clones recovered from the introduction of the editing plasmid (pAY6419) to PAO1^λIF^ to construct Δ*pqsA*. (**A**) Sequence of the mini-CRISPR array employed to construct *ΔpqsA* and that from eight false positive clones recovered from introduction of the editing plasmid (pAY6419) to PAO1^λIF^ to construct *ΔpqsA*. The mini-CRISPR array region from the eight recovered plasmids were subjected to sanger sequencing and aligned with the originally employed mini-CRISPR sequence. loss of the spacer and one repeat sequence was identified in two of the eight mini-CRISPRs (No.2 and No.8). (**B**) Mini-CRISPR sequence employed for *pqsA* targeting.

**Figure S11.**
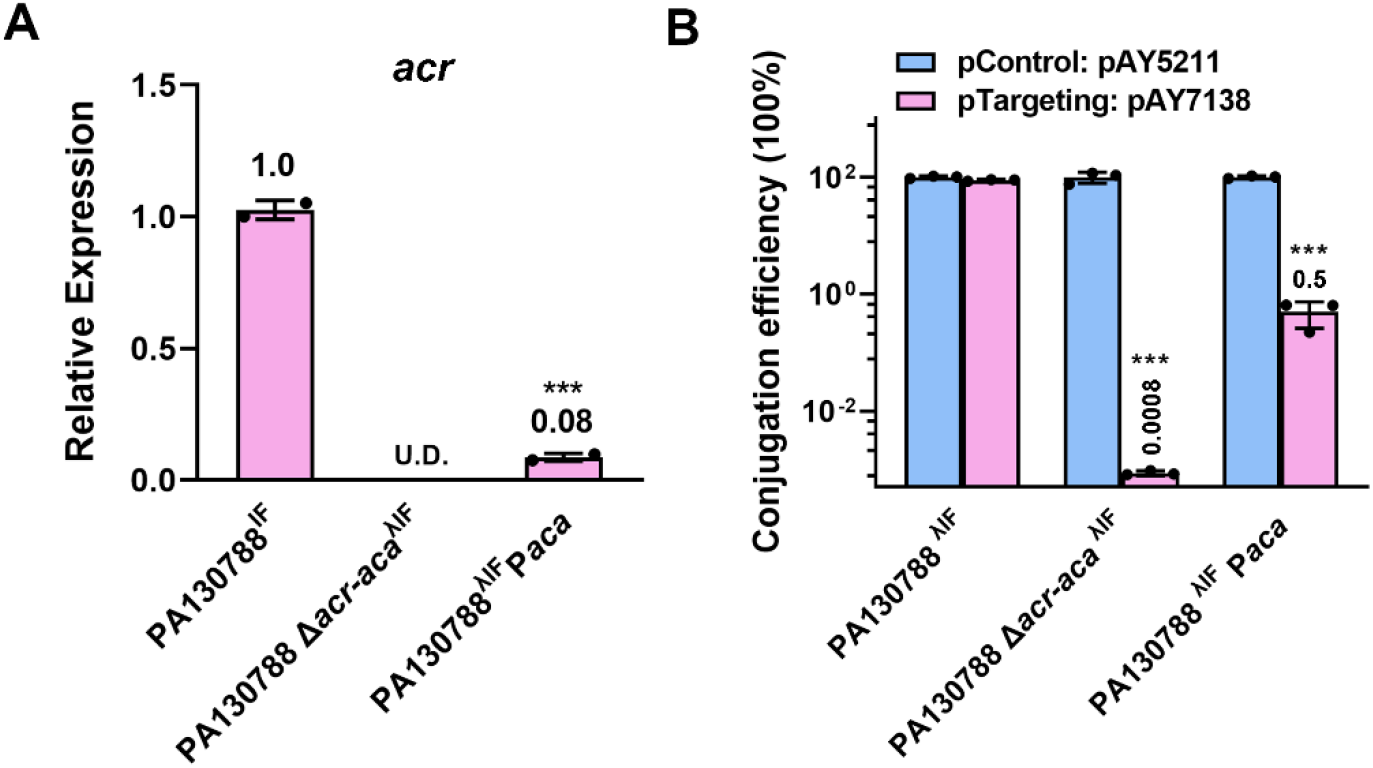
Overexpression of *aca* elevates the self-targeting capacity of PA130788. (**A**) Relative expression of *acr* in PA130788^λIF^, PA130788 Δ*acr-aca*^λIF^, and PA130788^λIF^ P*aca* strains. U.D.: Undetected. (**B**) DNA interference activity of the transferred I-F Cascade determined by the conjugation efficiency of transforming pAY7138 into the indicated host strains PA130788^λIF^, PA130788 Δ*acr-aca*^λIF^, and PA130788^λIF^ P*aca*. Data are the mean of two (A) or three (B) biological repeats and are expressed as mean ± S.D. Statistical significance is calculated based on student’s *t* test (***, *P* < 0.001).

**Table S1.**
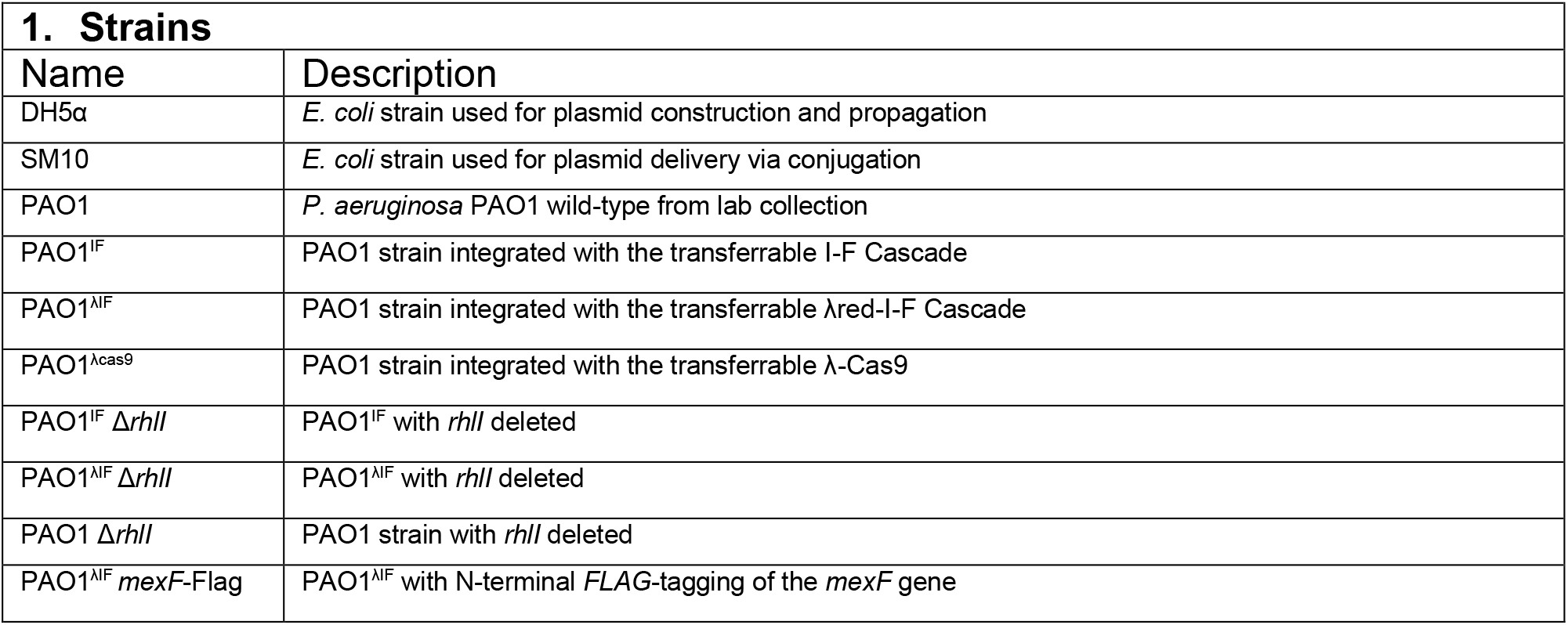
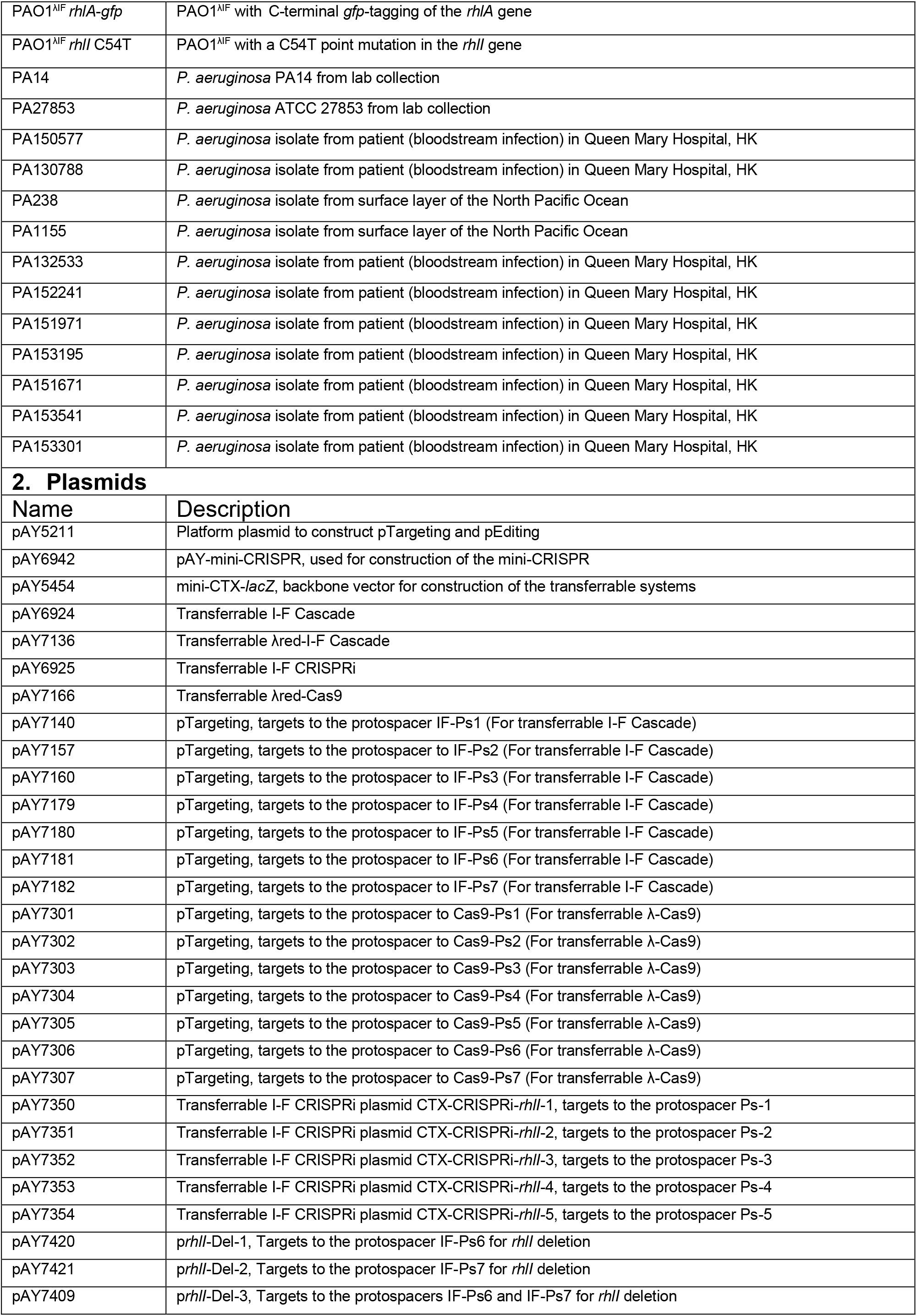
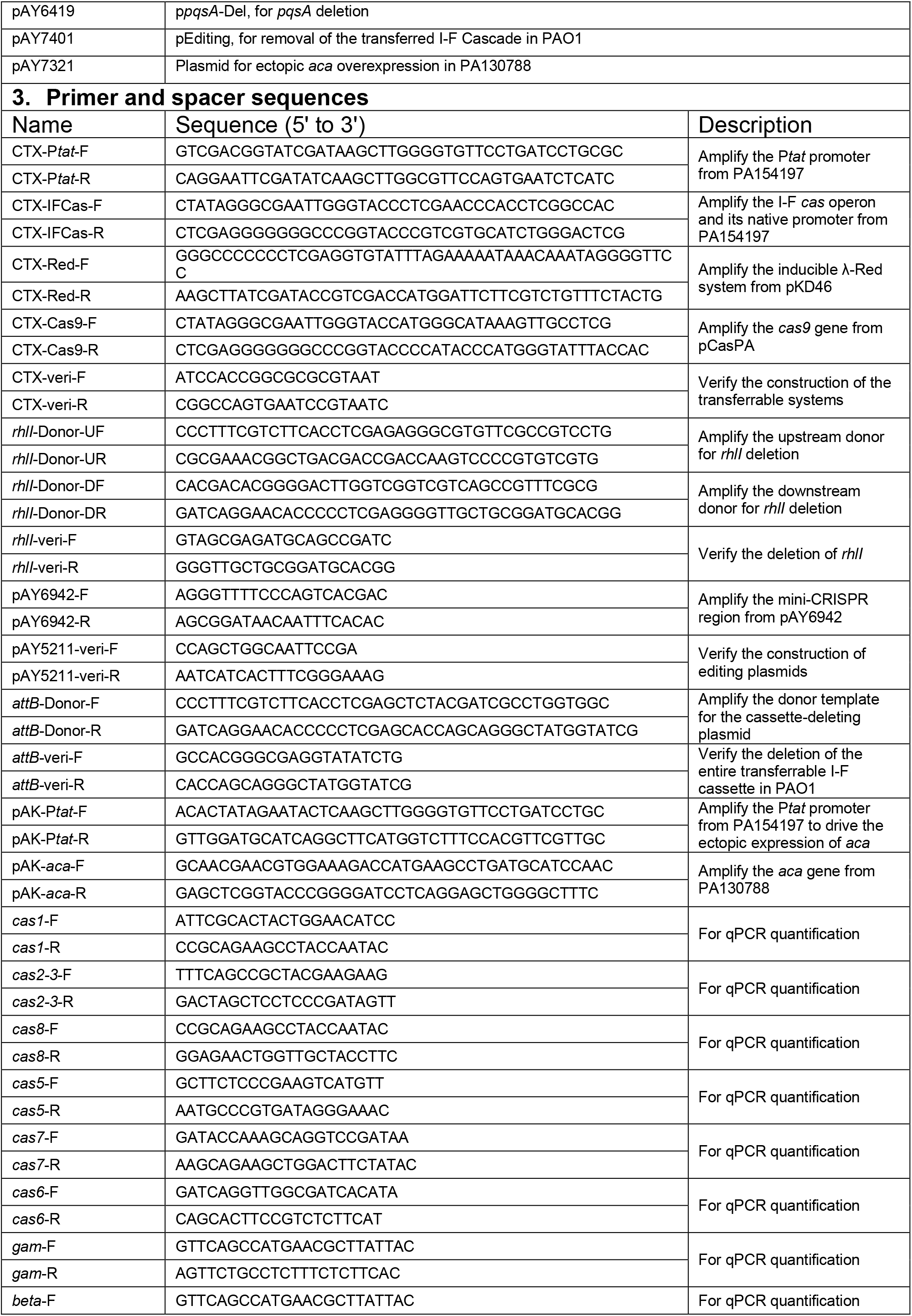
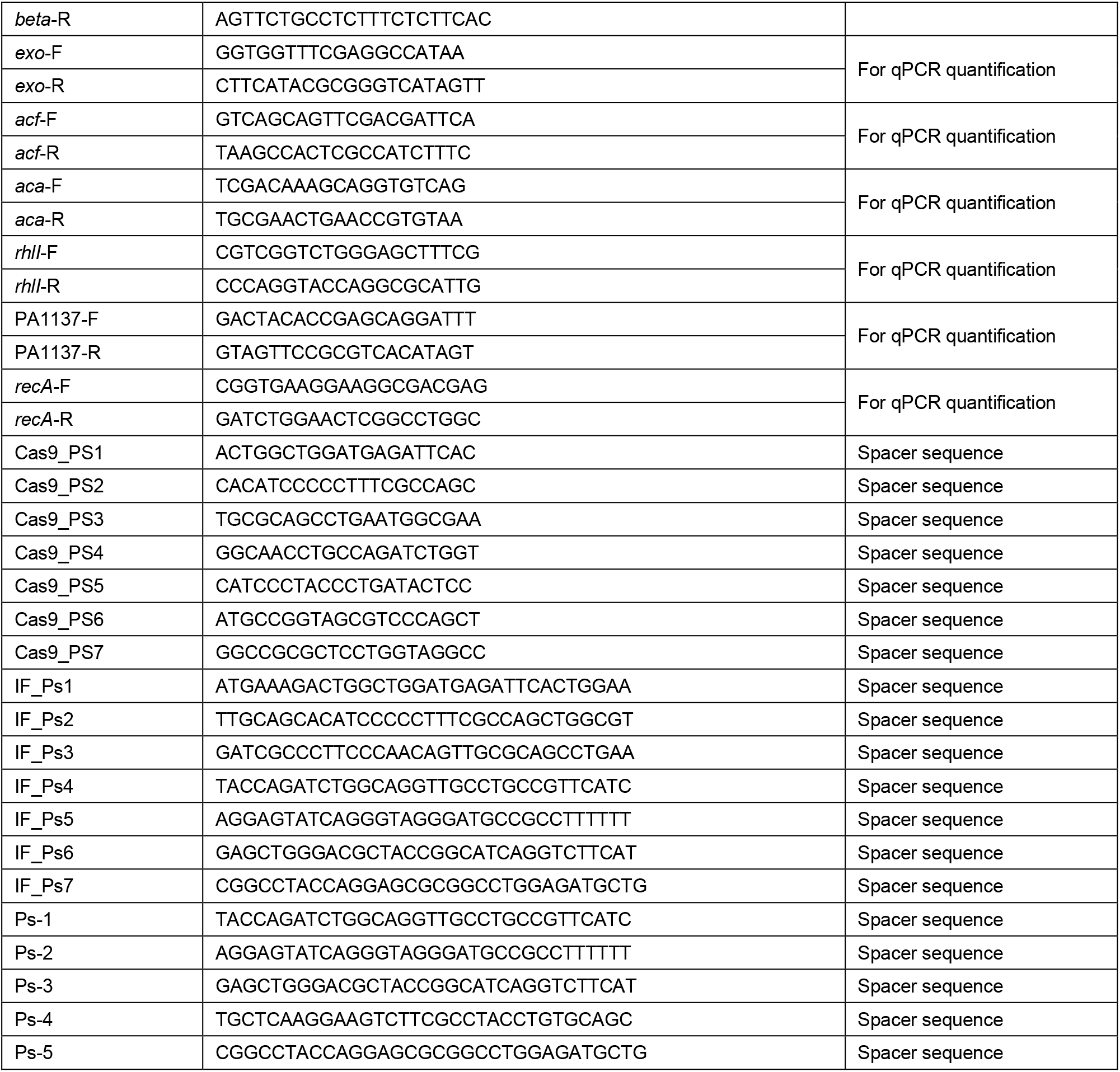
Bacterial strains, plasmids, primers, and spacers used in this study

**Table S2.**
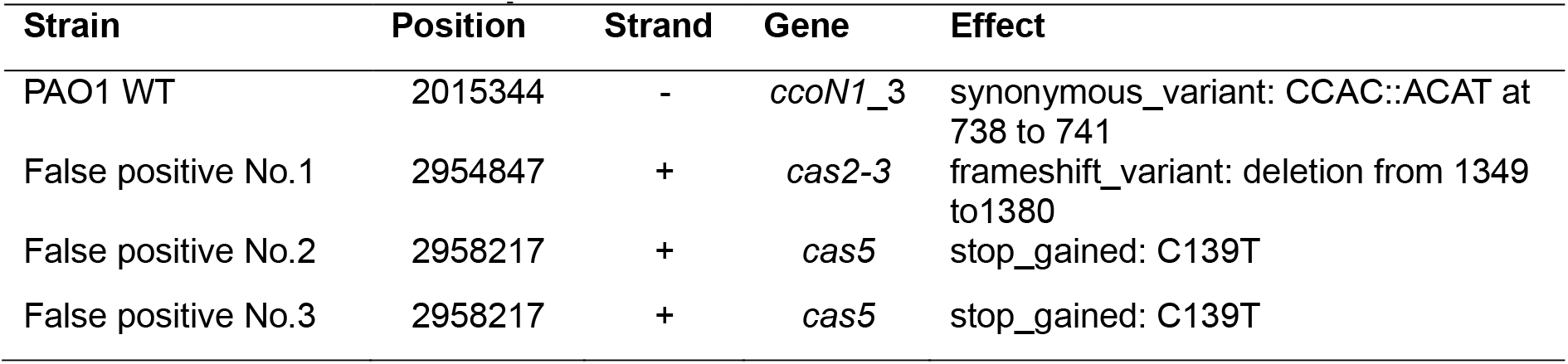
WGS revealed mutations in the false positive clones recovered from the introduction of *prhlI*-Del-1 into PAO1^λIF^

**Table S3.**
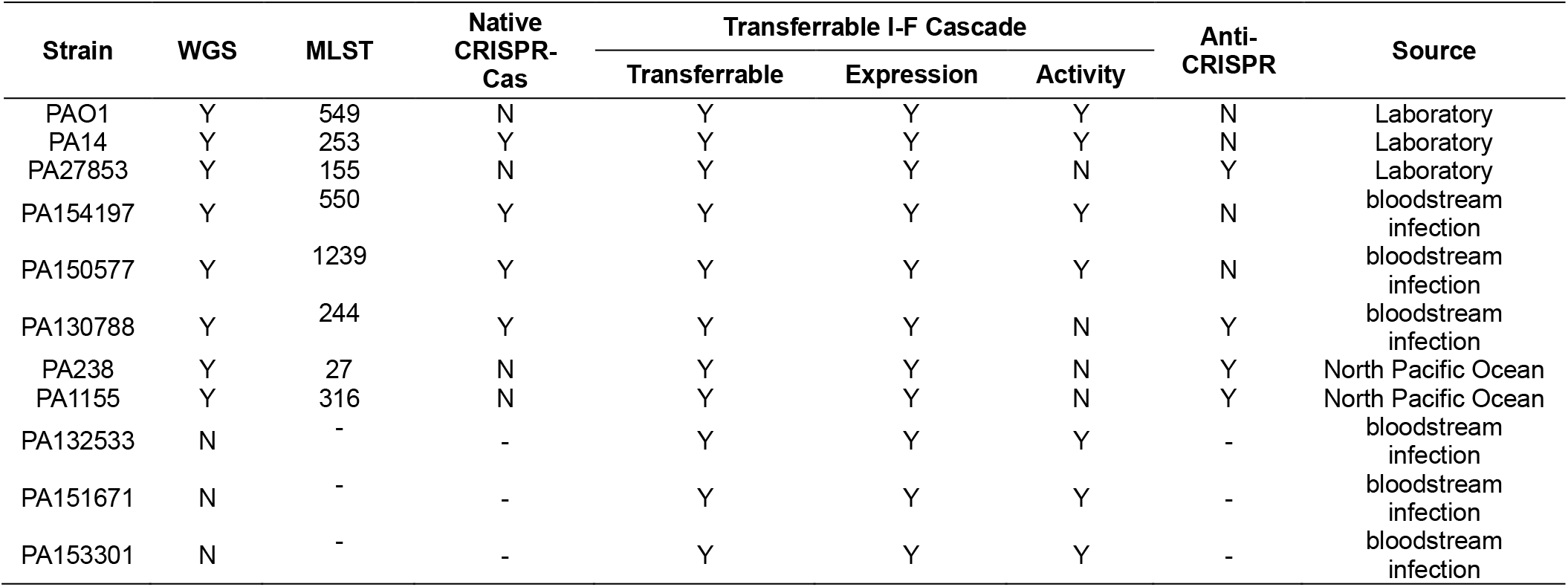
Summary of the *P. aeruginosa* strains in which the transferrable systems are applied

